# *In vivo* control of the ezrin/radixin/moesin protein ERM-1 in *C. elegans*

**DOI:** 10.1101/2020.01.08.898189

**Authors:** João J. Ramalho, Ophélie Nicolle, Grégoire Michaux, Mike Boxem

**Affiliations:** Division of Developmental Biology, Institute of Biodynamics and Biocomplexity, Department of Biology, Faculty of Science, Utrecht University, Padualaan 8, 3584 CH, Utrecht, The Netherlands; Univ Rennes, CNRS, IGDR (Institut de Génétique et de Développement de Rennes), UMR 6290, F-35000 Rennes, France; Present address: Laboratory of Biochemistry, Wageningen University & Research, Stippeneng 4, 6708 WE, Wageningen, The Netherlands

## Abstract

ERM proteins are conserved regulators of cortical membrane specialization, that function as membrane–actin linkers and molecular hubs. Activity of ERM proteins requires a conformational switch from an inactive cytoplasmic form into an active membrane- and actin-bound form, which is thought to be mediated by sequential PIP_2_-binding and phosphorylation of a conserved C-terminal threonine residue. Here, we use the single *C. elegans* ERM ortholog, ERM-1, to study the contribution of these regulatory events to ERM activity and tissue formation *in vivo*. Using CRISPR/Cas9-generated *erm-1* mutant alleles we demonstrate that PIP_2_-binding is critically required for ERM-1 function. In contrast, dynamic regulation of C-terminal T544 phosphorylation is not essential but modulates ERM-1 apical localization and dynamics in a tissue-specific manner, to control cortical actin organization and drive lumen formation in epithelial tubes. Our work highlights the dynamic nature of ERM protein regulation during tissue morphogenesis and the importance of C-terminal phosphorylation in fine-tuning ERM activity in a tissue-specific context.

## Introduction

Morphological and molecular specialization of defined regions at the cell cortex is critical for the development and function of most animal cell types. Formation of specialized cortical domains relies on local reorganization of membrane composition and the cortical cytoskeleton. Ezrin/Radixin/Moesin (ERM) proteins form an evolutionary conserved family that plays a major role in organizing the cell cortex and signaling (Fehon et al., 2010; McClatchey, 2014; Neisch and Fehon, 2011). For example, ERM family members promote the formation of microvilli at the apical surface of epithelial tissues, are required for lumen formation in tubular epithelia, and control the mechanical properties of the cell cortex in processes such as mitosis, cell migration, and immunological synapse formation in B and T cells (Kunda et al., 2008; McClatchey, 2014; Parameswaran and Gupta, 2013; Pelaseyed and Bretscher, 2018). To perform this wide range of functions, ERM proteins can physically link the plasma membrane with the actin cytoskeleton and orchestrate the assembly of a broad array of multiprotein complexes at the cell surface.

ERM proteins consist of an N-terminal band four-point-one/ezrin/radixin/moesin (FERM) domain that mediates binding to transmembrane- and membrane-associated proteins, a C-terminal tail that mediates actin binding, and a central α-helical linker region (Fehon et al., 2010; McClatchey, 2014). The activity of ERM proteins is regulated by a reversible intramolecular interaction. In the inactive closed conformation, extensive interactions between the FERM domain and the C-terminal tail mask the actin-binding site, membrane-binding sites, and protein interaction sites (Gary and Bretscher, 1995; Li et al., 2007; Magendantz et al., 1995; Pearson et al., 2000). The transition to a fully open and active conformation involves binding to the plasma membrane lipid phosphatidylinositol-(4,5) bisphosphate (PIP_2_) as well phosphorylation of a specific C-terminal threonine residue (T567 in ezrin, T564 in radixin and T558 in moesin) (Simons et al., 1998; Nakamura et al., 1999; Barret et al., 2000; Fievet et al., 2004; Coscoy et al., 2002; Yonemura et al., 2002; Hao et al., 2009; Roch et al., 2010). This transition is thought to occur in a multistep process, in which binding to PIP_2_ induces a partial conformational change that enables binding of a kinase and phosphorylation of the C-terminal threonine (Fievet et al., 2004; Hao et al., 2009; Bosk et al., 2011; Pelaseyed et al., 2017).

While binding to PIP_2_ appears to be absolutely essential for the activity of ERM proteins (Nakamura et al., 1999; Barret et al., 2000; Coscoy et al., 2002; Yonemura et al., 2002; Fievet et al., 2004; Hao et al., 2009; Roch et al., 2010), the role and contribution of phosphorylation of the C-terminal threonine residue to ERM protein activity *in vivo* is less clear. Numerous studies have indicated that phosphorylation corresponds with activation of ERM proteins, and phosphomimetic mutations induce the transition to an active membrane-bound state, promote actin binding, and induce formation of cortical actin-rich structures or cortical rigidity (Huang et al., 1999; Matsui et al., 1998; Nakamura et al., 1999; Oshiro et al., 1998; Simons et al., 1998; Gautreau et al., 2000; Polesello et al., 2002; Speck et al., 2003; Chambers and Bretscher, 2005; Charras et al., 2006; Carreno et al., 2008; Kunda et al., 2008; Bosk et al., 2011). However, more recent studies indicate that ERM proteins are not static membrane–actin linkers, but undergo rapid turnover at the membrane, accompanied by constant phosphorylation cycling of the C-terminal threonine (Viswanatha et al., 2012; Fritzsche et al., 2014; Parameswaran et al., 2011). Consistent with an essential role for phosphocycling, several studies in mammalian systems and *Drosophila* have shown that neither non-phosphorylatable nor phosphomimetic variants of ERM proteins can substitute for the wild-type protein (Viswanatha et al., 2012; Parameswaran et al., 2011; Polesello et al., 2002; Karagiosis and Ready, 2004). However, results in *Drosophila* are conflicting, as significant rescuing capacity has been reported for a non-phosphorylatable Moesin variant (Roch et al., 2010), and both rescuing and dominant-lethal effects have been reported for phosphomimetic Moesin variants (Polesello et al., 2002; Speck et al., 2003; Hipfner, 2004; Karagiosis and Ready, 2004). These studies highlight the complexity and importance of studying ERM regulation *in vivo*.

Here, we investigate the contributions of PIP_2_ binding and C-terminal threonine phosphorylation to the functioning of the single ERM protein ERM-1 in the nematode *Caenorhabditis elegans*. ERM-1 is highly similar in sequence and domain composition to ERM proteins in other organisms. Importantly, the residues critical for PIP_2_ binding as well as the C-terminal threonine residue are fully conserved (Fig. 1A). ERM-1 localizes to the apical surface of most polarized tissue types and is essential for apical membrane morphogenesis (Göbel et al., 2004; van Fürden et al., 2004). The loss of *erm-1* function causes early larval lethality with severe cystic defects in the intestine and excretory canals, two polarized tubular epithelia (Göbel et al., 2004). In the intestine, loss of *erm-1* causes constrictions, loss of microvilli, severe reduction in the levels of apical actin, and defects in the accumulation of junctional proteins (Bernadskaya et al., 2011; Göbel et al., 2004; van Fürden et al., 2004). The excretory canals are part of the *C. elegans* excretory system, thought to be involved in osmoregulation and secretion (Sundaram and Buechner, 2016). They are seamless tubes that extend on the lateral sides for nearly the entire length of the animal, with an apical domain forming the inner luminal surface, and a basal domain forming the outer wall of the tube (Sundaram and Buechner, 2016). ERM-1 localizes to the lumen-forming apical membrane, and controls the extension of the canal lumen in a dose-dependent manner (Khan et al., 2013; Göbel et al., 2004). In addition, loss of *erm-1* causes a highly vacuolated appearance of the canal cytoplasm (Khan et al., 2013).

**Figure 1.**
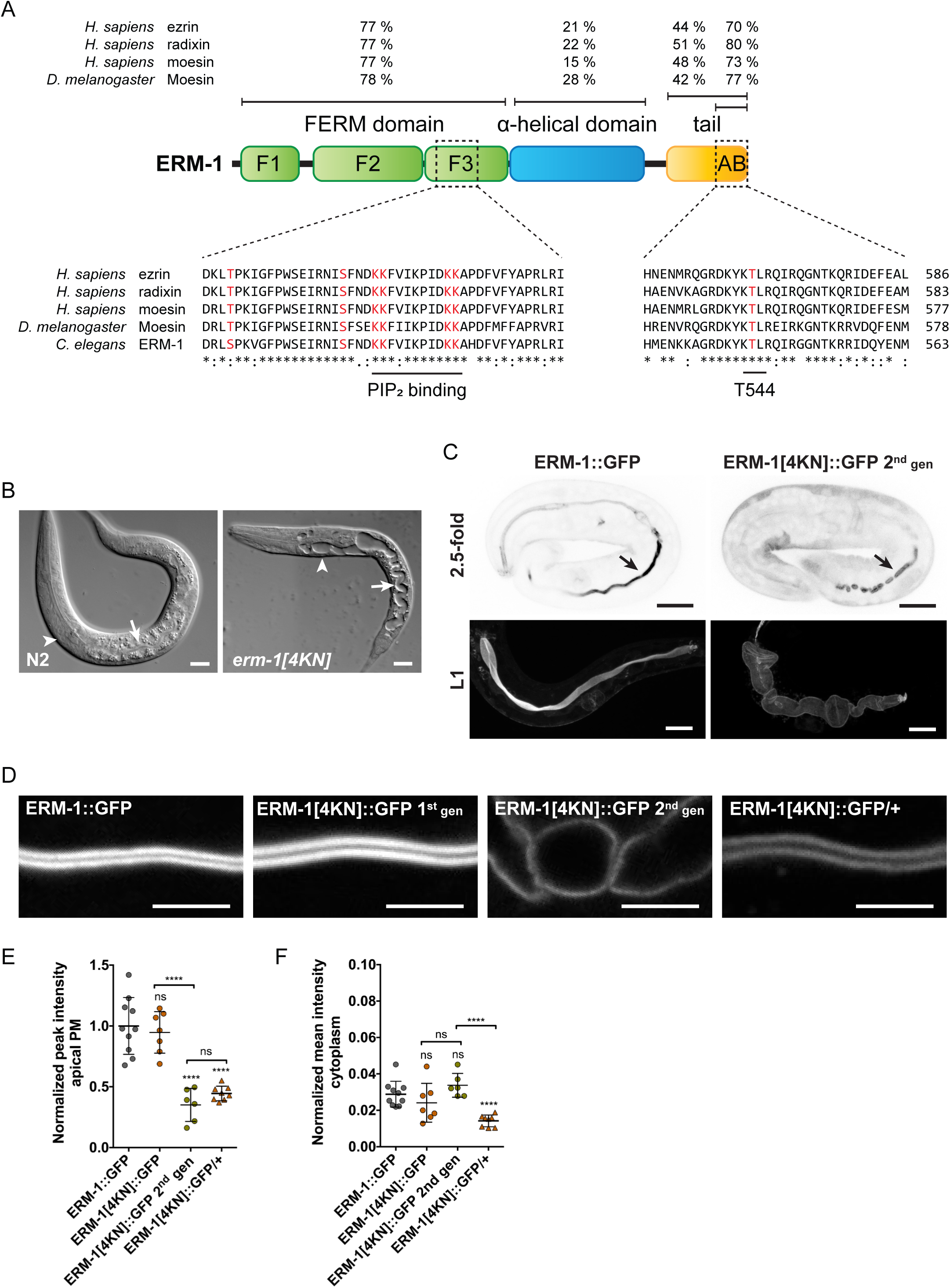
ERM-1 activity and efficient membrane targeting require binding to PIP_2_. (**A**) Schematic representation of the domain organization of ERM-1. F1-F3 correspond to the three structural modules making up the FERM domain, AB is the actin binding domain. (**B**) DIC microscopy images of N2 animals or PIP_2_-binding deficient *erm-1[4KN]* mutants. Arrowheads point to the excretory canal, and arrows to the intestinal lumen. (**C**) Distribution of ERM-1::GFP and ERM-1[4KN]::GFP second generation homozygotes in late embryos and L1 larvae. Arrow points to the intestinal apical membrane. (**D**) Distribution of GFP-tagged ERM-1 variants in the intestine of L1 larvae with indicated genotypes. Images were acquired and displayed with the same settings (C,D). Images correspond to single planes (B,D), or maximum intensity projections (C). In this and all other figures anterior is to the left, and unless otherwise indicated imaging was done with spinning-disk confocal microscopy. Scale bars, 10 µm. (**E,F**) Plots of GFP mean intensity quantified at the apical membrane (E) or cytoplasm (F) in intestines of L1 larvae as shown in (D). Values are normalized to the mean intensity of ERM-1::GFP at the apical membrane. Each symbol represents the average of 6 measurements per animal for the apical membrane, and 3 for the cytoplasm; n≥6. Unless indicated otherwise by a connecting line, statistical comparisons are with ERM-1::GFP. Unless otherwise indicated data in this and all other figures are presented as mean ± SD, and analyzed with unpaired t-test; ns: p-value > 0.05; *: p-value < 0.05; **: p-value < 0.01; ***: p-value < 0.001; ****: p-value < 0.0001.

We used CRISPR/Cas9 to endogenously engineer the *erm-1* locus to express ERM-1 variants that are incapable of binding to PIP_2_, cannot be phosphorylated, or mimic phosphorylation. As in other systems, PIP_2_ binding was essential for the functioning of ERM-1. In contrast, animals expressing only non-phosphorylatable or phosphomimetic ERM-1 protein were viable, demonstrating unequivocally that C-terminal threonine phosphorylation is not strictly required for ERM activity in *C. elegans.* Nevertheless, phosphorylation contributed to multiple aspects of ERM-1 function, including localization, mobility, and ability to organize an apical actin network. Effects caused by the phosphorylation mutants were often tissue-specific, highlighting a surprising versatility for C-terminal phosphorylation in controlling ERM-1. Finally, in support of an essential role for phosphorylation cycling, most of the defects we observed were highly similar between non-phosphorylatable and phosphomimetic variants.

## Results

### Lipid binding is essential for the activity of ERM-1

Binding to PIP_2_ has been shown to be essential for the membrane localization and activity of ERM proteins in multiple systems, including mammalian tissue culture cells and *Drosophila* (Nakamura et al., 1999; Barret et al., 2000; Coscoy et al., 2002; Yonemura et al., 2002; Fievet et al., 2004; Hao et al., 2009; Roch et al., 2010). We first sought to confirm that binding to PIP_2_ is critical for the functioning of *C. elegans* ERM-1 as well. Mutation of two pairs of lysine residues within a plasma-binding patch region to asparagines has been shown to virtually abolish membrane localization of ezrin in tissue culture cells (Barret et al., 2000; Fievet et al., 2004; Ben-Aissa et al., 2012), and fly Moesin carrying K–N mutations is unable to provide rescuing activity (Roch et al., 2010). We used CRISPR/Cas9 to change the corresponding lysine doublets in *C. elegans* ERM-1 to asparagine (K254/255N and K263/264N, from here on referred to as 4KN) (Fig. 1A). The first homozygous generation of *erm-1(mib11*[4KN]) is viable, while >95% of their offspring arrest during larval development, mostly as L1, with cysts in the lumens of the intestine and excretory canals (Fig. 1B). Both the maternal effect lethality and the cystic luminal phenotypes have been described for the putative *erm-1(tm667*) null allele (Göbel et al., 2004). In addition, first generation homozygous *erm-1(mib11[4KN])* animals die prematurely in adulthood, as has been observed for *erm-1(tm677)*. Thus, *erm-1(mib11[4KN])* behaves similar to a putative null allele.

To examine whether the ERM-1[4KN] mutation prevents membrane association, we generated endogenous COOH-terminal fusions of eGFP (referred to as GFP) with wild-type ERM-1 and ERM-1[4KN], using CRISPR/Cas9. We observed expression of ERM-1::GFP in all developmental stages (Fig. 1C,D; Fig. S1A,G). In early embryogenesis, ERM-1::GFP localized to the entire plasma membrane as well as the cytoplasm (Fig. S1A). As morphogenesis initiates, ERM-1::GFP was primarily detected at the apical surface of epithelial tissues and primordial germ cells (PGCs) (Fig. S1A). In larval stages, we observed apical localization of ERM-1::GFP in the intestine, seam cells, and excretory canals (Fig. 1D; Fig. S1G). In the syncytial germline, ERM-1 was associated with the entire plasma membrane, but enriched at the apical domain (Fig. S1G). We also observed ERM-1 in other tissues including P cells, vulva, uterus, spermatheca, and several neurons (data not shown).

Addition of N- or C-terminal tags to ERM proteins has been suggested to interfere with the intramolecular interaction between the FERM-domain and C-terminus (Chambers and Bretscher, 2005; Viswanatha et al., 2012). Consistent with this, we observed that *erm-1::GFP* animals have a reduced brood size and incomplete outgrowth of the excretory canals (Figure S1B-E). This effect is likely not due to constitutive ERM-1 activity, as excretory canal defects are absent in *erm-1::GFP/+* heterozygotes (Fig. S1. D,E). However, C-terminal GFP fusions have been used extensively to characterize the distribution of ERM proteins (Coscoy et al., 2002; Göbel et al., 2004; Roch et al., 2010; Viswanatha et al., 2012; Garbett and Bretscher, 2012; Khan et al., 2013; Babich and Di Sole, 2015), and the localization of our endogenous ERM-1::GFP fusion mimicked previous reports of ERM-1 localization in *C. elegans* (Göbel et al., 2004; van Fürden et al., 2004; Bernadskaya et al., 2011; Khan et al., 2013). ERM-1::GFP therefore appears to accurately reflect the localization of the endogenous protein.

Similar to mammalian ezrin and *Drosophila* Moesin PIP_2_ binding mutants (Barret et al., 2000; Roch et al., 2010), ERM-1[4KN]::GFP failed to localize to the plasma membrane in PGCs and seam cells of second generation homozygous *erm-1[4KN]::GFP* L1 larvae (Fig. S1G). We also detected a severe reduction of apical ERM-1[4KN]::GFP in the intestine (Fig. 1C–F). Surprisingly, ERM-1[4KN]::GFP appeared to still be restricted to the apical domain of the excretory canal (Fig. S1G). Mammalian and *Drosophila* ERM PIP_2_ binding mutants have been reported to relocalize to the cytoplasm. We observed clear cytoplasmic relocalization in the PGCs and germline during larval stages, but not in the intestine or excretory canal (Fig. 1D,F; Fig. S1G), indicating that cytoplasmic ERM-1 or ERM-1[4KN] may not be stable in these cell types.

To further characterize the effect of the 4KN mutation on ERM-1 membrane targeting we analyzed the distribution of ERM-1[4KN]::GFP in heterozygous *erm-1[4KN]::GFP/+* animals. While ERM-1[4KN]::GFP still failed to localize to the membrane in PGCs and seam cells, in the intestine, excretory canal, and larval germline we observed clear apical localization of ERM-1[4KN]::GFP at the apical membrane domain (Fig. 1D; Fig. S1G). Interestingly, in first generation homozygous animals, apical levels of ERM-1[4KN]::GFP were comparable to wild-type ERM-1::GFP (Fig. 1D-G). These animals have low levels of maternally contributed wild-type ERM-1. Thus, it appear that wild-type ERM-1 is able to support apical recruitment of ERM-1[4KN], though not in all tissues. These results contrast with observations in *Drosophila* and mammalian tissue culture cells where C-terminally tagged ERM 4KN mutants do not localize to the plasma membrane, even in the presence of wild-type ezrin (Babich and Di Sole, 2015; Barret et al., 2000; Fievet et al., 2004; Hao et al., 2009; Roch et al., 2010).

Based on the severe loss-of-function phenotype caused by *erm-1(mib11[4KN])* and the altered membrane association observed for ERM-1[4KN]::GFP, we conclude that membrane localization, presumably through binding to PIP_2_, is critical for activity of *C. elegans* ERM-1, as it is for ERM proteins in other organisms. However, the presence of wild-type ERM-1 protein is able to compensate for the loss of membrane-binding activity of the ERM-1[4KN] mutant at least in some tissues.

### Phosphorylation of T544 is not essential for ERM-1 functioning

We next investigated the contribution of phosphorylation of the conserved C-terminal regulatory threonine residue to ERM-1 functioning (Fig. 1A). We determined the extent of phosphorylation of ERM-1 using an antibody that specifically recognizes C-terminally phosphorylated ERM proteins (pERM). We observed extensive pERM staining throughout development, including at apical membranes in the intestine, seam cells, and excretory canals, and throughout the entire plasma membrane in the adult germline and oocytes (Fig. 2A). We did not observe pERM staining in animals in which T544 was replaced with an alanine (see below), indicating that the pERM antibody is specific for the phosphorylated form of ERM-1 (Fig. 2B). Thus, ERM-1 is broadly phosphorylated on the conserved C-terminal threonine residue.

**Figure 2.**
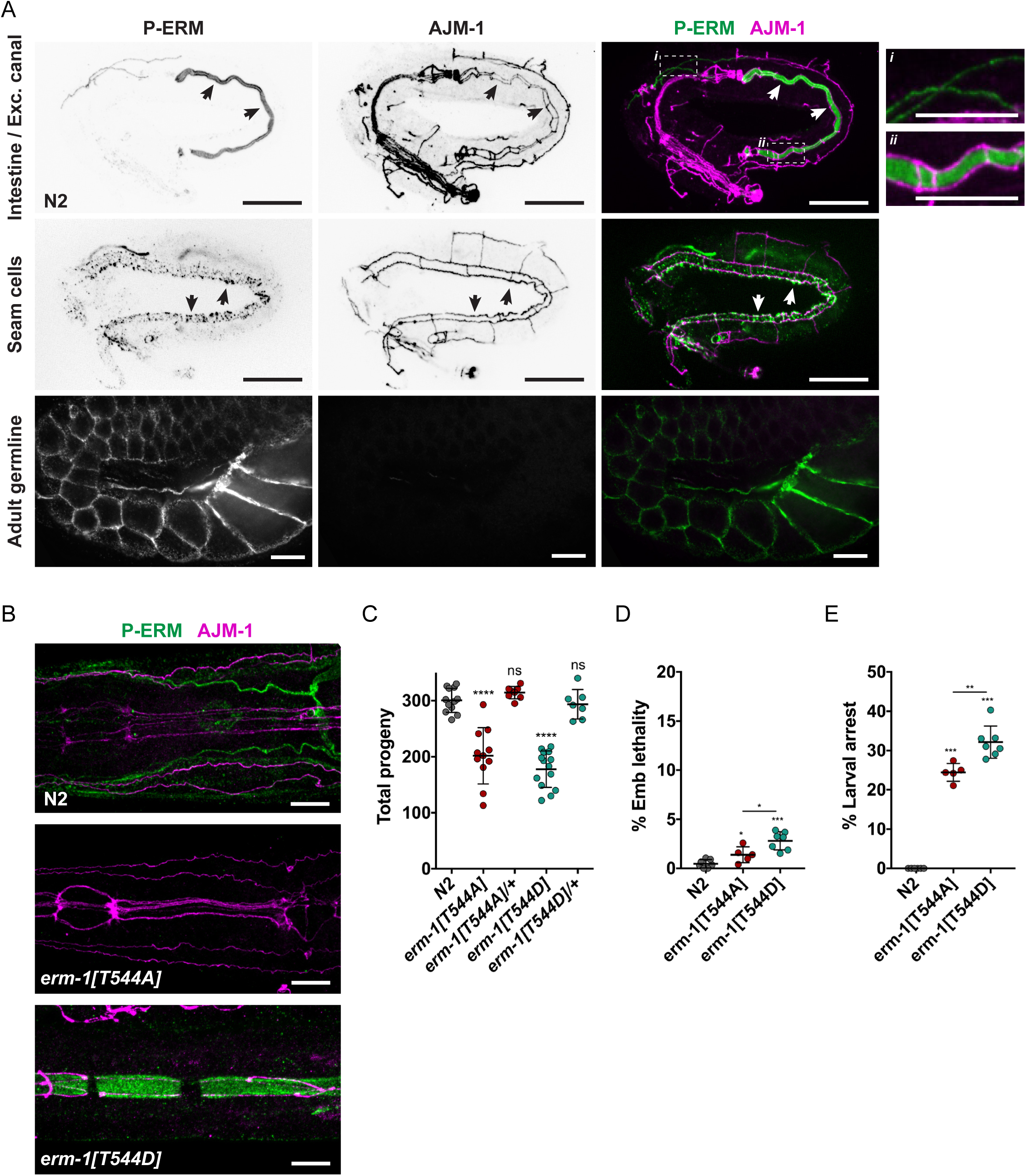
T544 is phosphorylated *in vivo* but phosphorylation is not essential for ERM-1 activity. (**A**) Distribution of T544-phosphorylated ERM-1 (pERM) and junctional marker AJM-1 in different embryonic (top and middle) and adult (bottom) tissues assessed by antibody staining of fixed wild type animals. Arrows point to the intestinal lumen (top) or seam cells (middle). Far right panels are enlarged views of the corresponding highlighted regions. (**B**) Distribution of pERM and AJM-1 in the pharynx or intestine of fixed adult N2 animals, or mutants homozygous for non-phosphorylated (T544A) or phosphomimetic (T544D) *erm-1* alleles. Images are maximum intensity projections. Scale bars: 10 µm; blow-ups in (A) 5 µm. (**C-E**) Quantification of total hatched progeny (C), as well as percentage of embryonic (D) and larval lethality (E) per animal. Each symbol represents the progeny of an individual animal; n≥5. Unless indicated otherwise by a connecting line, statistical comparisons are with N2.

To assess the importance of T544 phosphorylation *in vivo*, we used CRISPR/Cas9 to generate *erm-1* mutants in which T544 is replaced with an alanine (T544A) or aspartic acid (T544D), to mimic the non-phosphorylated and phosphorylated states respectively (Fig. 1A). We detected pERM staining in intestinal cells of *erm-1[T544D]* mutants, suggesting that this variant indeed mimics the phosphorylated form (Fig. 2B). Surprisingly, both homozygous non-phosphorylatable *erm-1[T544A]* and phosphorylation mimicking *erm-1[T544D]* mutants are viable, demonstrating that T544 phosphorylation is not essential for ERM-1 activity in *C. elegans*. Nevertheless, both mutants have a reduced brood size, and increased embryonic and larval lethality compared to wild type animals (Fig. 2C–E). These defects were not observed in heterozygous *erm-1[T544A]/+* and *erm-1[T544D]/+* animals, indicating that neither mutation exerts a dominant effect. Finally, consistent with the broad expression pattern of ERM-1, we observed a variety of partially penetrant phenotypes, including small, dumpy, tail defects, protruding vulva, exploded through vulva, uncoordinated, and clear. Together, these results demonstrate that the C-terminal phosphorylation site is important, but not required, for ERM-1 function in *C. elegans*.

### T544 phosphorylation contributes to lumen formation in tubular epithelia

To better understand the defects caused by mutation of T544, we investigated the effects on the intestine and excretory canal, tissues in which the role of ERM-1 has been extensively documented (Göbel et al., 2004; Khan et al., 2013; van Fürden et al., 2004). We examined the appearance and extension of the canal tubes using a COOH-terminal VHA-5∷GFP fusion protein, which localizes to the apical membrane and canaliculi (Liégeois et al., 2006, 2007). In both *erm-1[T544A]* and *erm-1[T544D]* animals, we observed severe canal extension defects, as well as cystic canals and widened canal lumens (Fig. 3A,B). In most animals, the appearance of the canals was indistinguishable between *erm-1[T544A]* and *erm-1[T544D]* mutations. However, severely affected canals were slightly more abundant in *erm-1[T544D]* mutants (Fig. 3B,C). In accordance with a non-dominant effect of both mutations, canal defects were absent from heterozygous *erm-1[T544A]/+* and *erm-1[T544D]/+* animals (Fig. 3B,C). We also examined canal morphology by transmission electron microscopy, and observed widened lumens and canals in *erm-1[T544A]* and *erm-1[T544D]* animals (Fig. 3D). Overexpression of ERM-1 leads to lumen formation defects in the excretory canal, which can be rescued by depletion of the ERM-1 binding-partner aquaporin AQP-8, (Khan et al., 2013). As phosphorylation mimicking ERM variants are widely assumed to have constitutive activity, we assessed whether cyst formation in T544 mutants also requires AQP-8 activity. We depleted *aqp-8* by RNAi in *erm-1[T544A]* and *erm-1[T544D]* mutants and analyzed excretory canal morphology and outgrowth. As previously observed (Khan et al., 2013), wild-type animals treated with *aqp-8* RNAi showed normal morphology and a mild canal outgrowth phenotype (Fig. 3E). In both *erm-1[T544A]* and *erm-1[T544D]* mutants, depletion of *aqp-8* enhanced the severity of canal morphology and outgrowth defects (Fig. 3E–G). These observations suggest that defects in *erm-1[T544D]* are likely not due to increased ERM-1 activity, and that ERM-1 requires both T544 phosphorylation/dephosphorylation and AQP-8 activity to promote lumen formation in the excretory canal.

**Figure 3.**
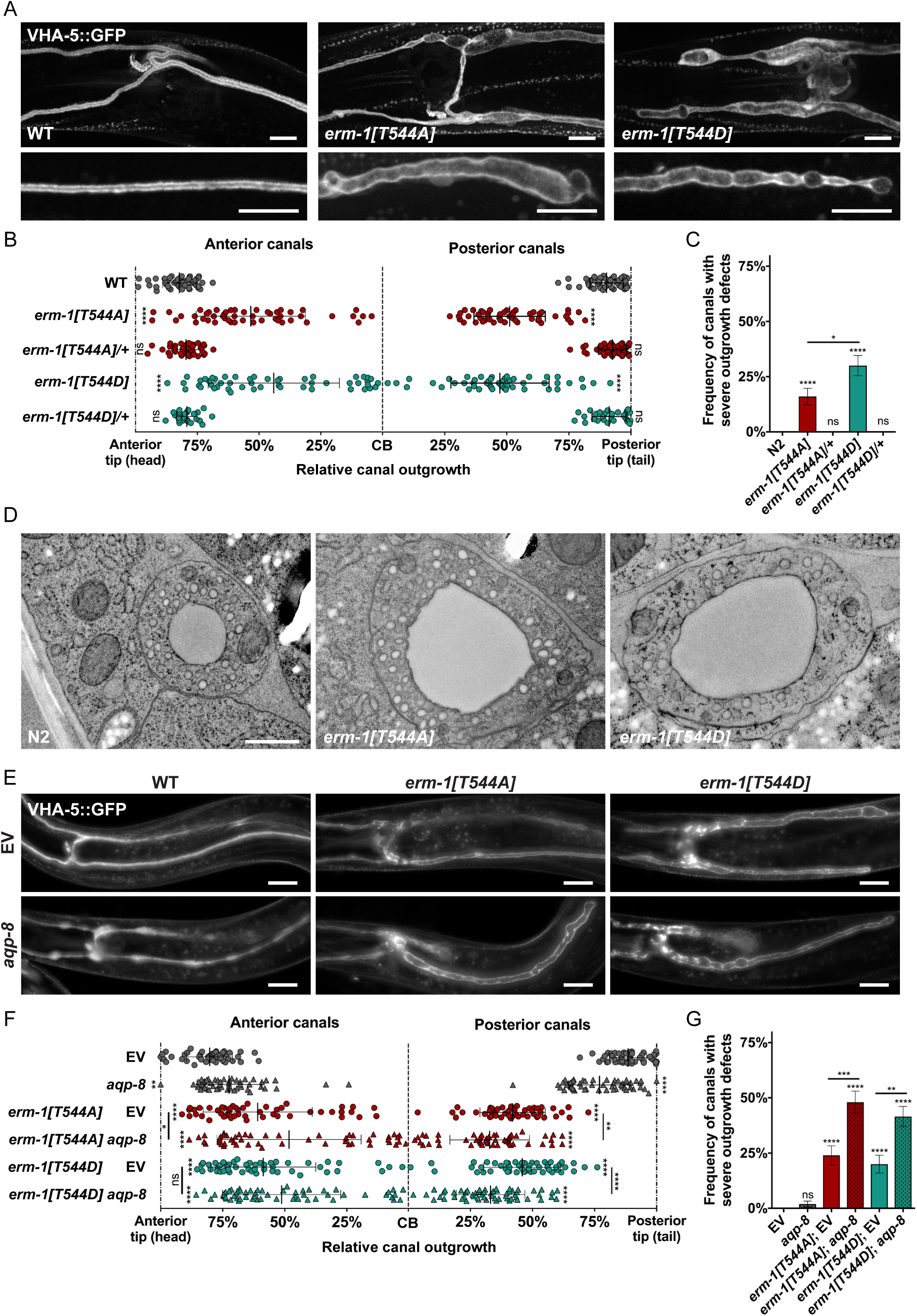
Dynamic regulation of ERM-1 T544 phosphorylation is required for lumen formation in the excretory canal in parallel to ERM-1/AQP-8 interaction. (**A**) Excretory canal lumen morphology in L4 animals visualized by a VHA-5::GFP transgene at the cell body region (top) or along the posterior canals (bottom). (**B**) Quantification of relative excretory canal outgrowth from the cell body to the anterior or posterior tips in L4 animals. All four canal branches were measured per animal, and each data point represents one branch; n≥14. (**C**) Frequency of anterior and posterior canals from (B) extending less than 35% of the distance between cell body and tips. (**D**) Transmission electron microscopy images of the excretory canal in L4 animals. (**E**) Excretory canal lumen morphology in L4 animals fed empty vector (EV) or *aqp-8* RNAi, visualized by a VHA-5::GFP transgene using epifluorescence microscopy. (**F,G**) Quantification of relative canal outgrowth and percentage of canals with severe defects as described above in L4 animals fed EV or *aqp-8* RNAi; n≥25. Images are maximum intensity projections. Scale bars: (A,E) 10 µm, (D) 200 nm. Unless indicated otherwise by a connecting line, statistical comparisons are with N2 (C) or EV (G).

We next investigated intestinal lumen formation by staining T544 mutant embryos with an antibody directed against the junctional protein AJM-1 (Francis and Waterston, 1991). In wild-type embryos, starting from the bean stage, junction bands on opposing sides of the expanding lumen visibly separate (Fig. 4A). In contrast, in both *erm-1[T544A]* and *erm-1[T544D]* mutants we did not detect clear separation of opposing junctions until the comma stage (Fig. 4A). Between comma and 2-fold stages, junction separation became visible but the distance was reduced compared to wild type animals, and constrictions that lack separation were visible along the length of the intestinal epithelium (Fig. 4A). At the 2-fold stage, *erm-1[T544D]* animals showed an almost complete separation of junctions throughout the epithelium (Fig 4A). In addition, we observed an apparent expansion or ectopic accumulation of junctional material (Fig. 4A). In contrast, in *erm-1[T544A]* embryos, junction constrictions were still often detected at the 2-fold stage (Fig. 4B).

**Figure 4.**
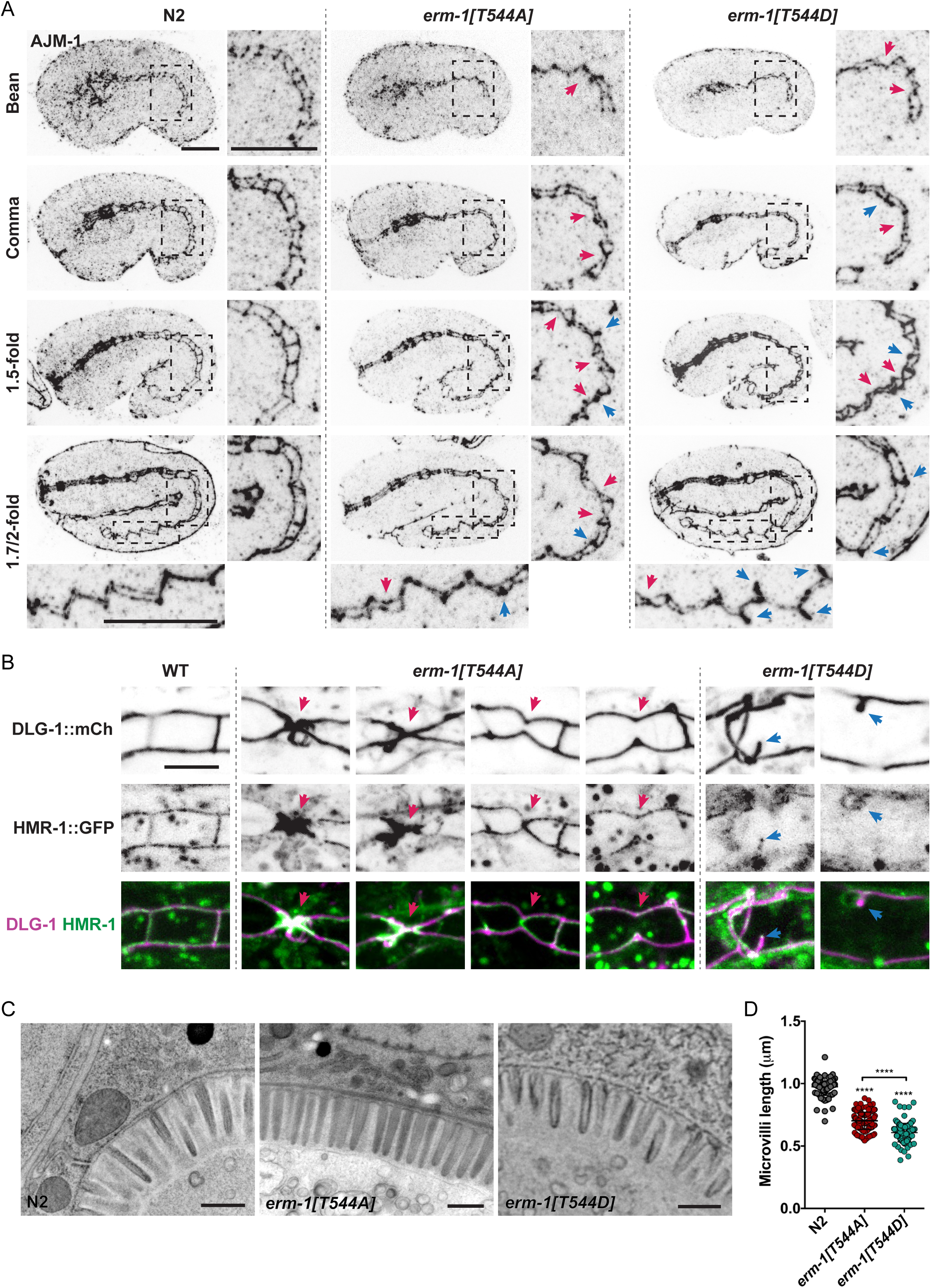
Positioning of cell junctions in intestinal cells requires dynamic regulation of ERM-1 T544 phosphorylation. (**A**) Junction organization visualized by AJM-1 antibody staining of fixed embryos at different embryonic stages. Images to the right of each panel and bottom of the last panel are enlarged views of the highlighted regions. Blue arrows highlight expansion or ectopic accumulation of junction material, and red arrows point to partial and full constrictions. Images are maximum intensity projections. (**B**) Junction organization in live L1 larvae expressing DLG-1::mCherry and HMR-1::GFP knock-ins. (**C**) Transmission electron microscopy images of the intestinal microvilli in L4 animals. (**D**) Quantification of microvilli length. Eeach symbol corresponds to a single microvillus, n≥4 animals (total microvilli quantified WT = 88, *erm-1[T544A]* = 168, *erm-1[T544D]* = 135). Unless indicated otherwise by a connecting line, statistical comparisons are with N2. Scale bars: (A) 10 µm; (B) 5 µm; (C) 500 nm.

We further characterized the junction defects of T544 ERM-1 mutants in living animals during larval stages, using a strain that expresses two endogenously tagged *C. elegans* apical junction proteins. DLG-1::mCherry marks the basal DLG-1/AJM-1 Complex (DAC), while HMR-1::GFP/E-cadherin is part of the more apical cadherin-catenin complex (CCC) (Pasti and Labouesse, 2014). We rarely observed full constrictions in *erm-1[T544D]* at the first larval stage (L1), although the junctions had a wavy appearance, and formed occasional aggregates as well as ectopic ring-like structures (Fig. 4B). Junctions in *erm-1[T544A]* animals had a higher frequency of partial or full constrictions (Fig. 4B). However, we did not detect significant junction defects in either mutant at subsequent larval stages (data not shown).

The lumen formation defects we observed in early intestinal development are similar to those reported for knockdown of *erm-1* by RNAi, which include reduced or complete failure in junction separation (van Fürden et al., 2004). The mild defects in intestines of L1 larvae and the observed recovery at later stages are however in stark contrast with previous descriptions of *erm-1* mutants or RNAi (Göbel et al., 2004; van Fürden et al., 2004), suggesting that T544 phosphorylation is important for ERM-1 activity during intestinal lumen formation, but is not strictly required.

ERM proteins in mammals, *Drosophila*, and *C. elegans*, are all critical for the formation of microvilli (Bonilha et al., 1999; Speck et al., 2003; Göbel et al., 2004; Saotome et al., 2004; Karagiosis and Ready, 2004; Bonilha et al., 2006; Casaletto et al., 2011), and C-terminal phosphorylation is thought to be essential for ERM proteins to support microvilli formation (Chen et al., 1995; Kondo et al., 1997; Gautreau et al., 2000; Pelaseyed et al., 2017; Viswanatha et al., 2012). We therefore examined the formation of intestinal microvilli in wild type animals, as well as animals expressing ERM-1[T544A] or ERM-1[T544D], by electron microscopy. Surprisingly, microvilli were still present in both mutant backgrounds at different larval stages (Fig. 4C), although their length was reduced in L4 larvae of both *erm-1[T544]* mutants, and more noticeably in *erm-1[T544D]* (Fig. 4D). Collectively, our results show that ERM-1 phosphorylation plays a major role in lumen formation during early embryogenesis, but is not essential for formation of a functional intestine or microvilli formation.

### Dynamic ERM-1 T544 phosphorylation contributes to molecular specialization of the apical domain

We next addressed whether ERM-1 T544 mutations perturb the functional specialization of the intestinal apical membrane. To this end, we introduced the *erm-1* T544 alleles in a strain overexpressing reporters for the peptide transporter PEPT-1::DsRed, which marks the apical membrane, and the small GTPase GFP::RAB-11, which marks apically enriched recycling endosomes (Winter et al., 2012). These proteins have been well characterized in *C. elegans*, and are known to have important functions in intestinal development and homeostasis across species (Spanier, 2014; Welz et al., 2014). In L1 larvae, we observed a dramatic reduction in the levels of PEPT-1::DsRed at the apical membrane in both *erm-1[T544A]* and *erm-1[T544D]* mutants (Fig. 5A, C). Both mutants also showed a reduction in GFP::RAB-11 enrichment near the apical plasma, although the effect was more pronounced in *erm-1[T544A]* (Fig. 5B, D). Thus, the balance between phosphorylated and non-phosphorylated ERM-1 forms contributes to the molecular specialization of the apical domain.

**Figure 5.**
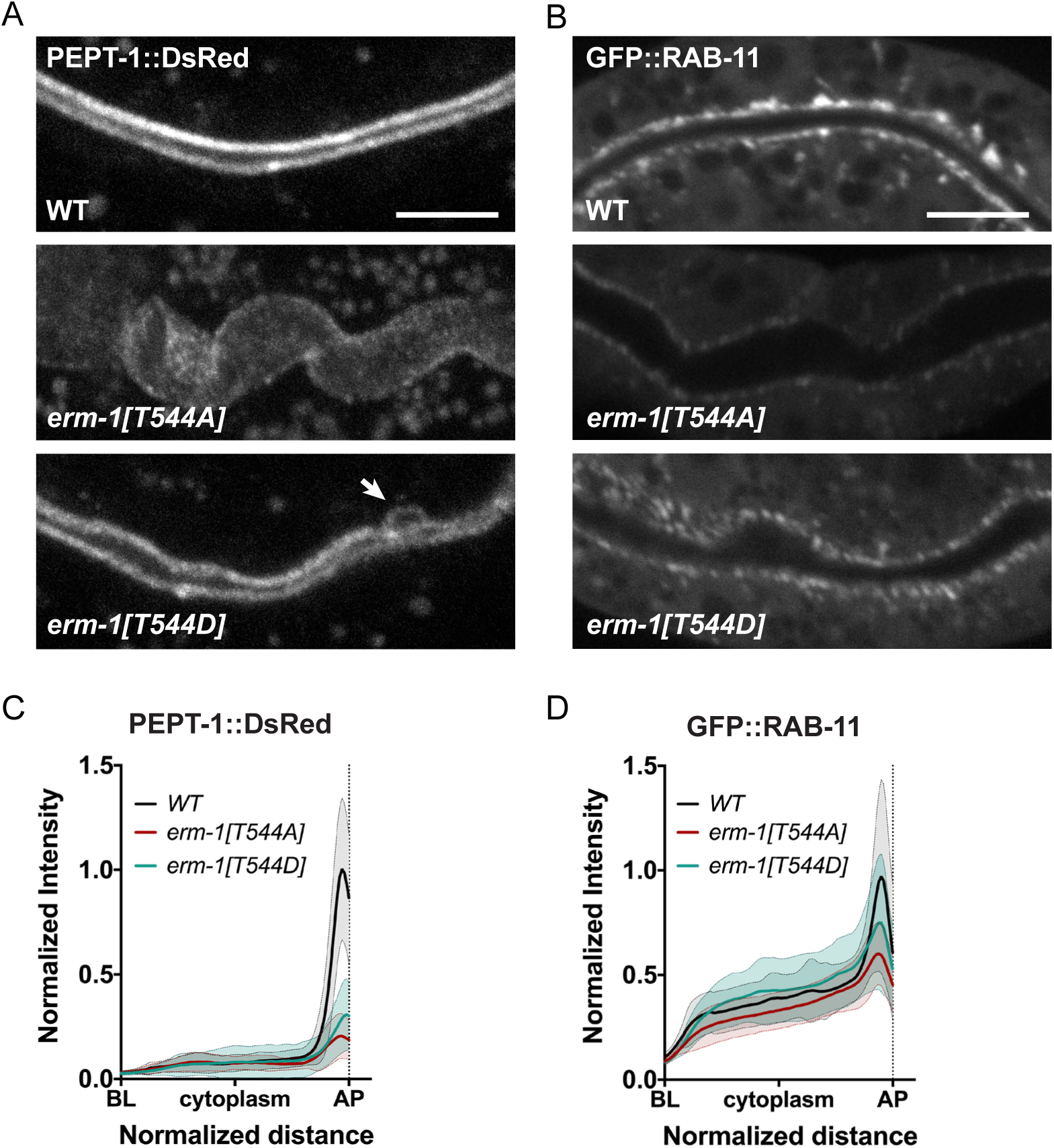
Dynamic T544 phosphorylation is important for molecular specialization of the apical domain. (**A,B**) Intestinal distribution of PEPT-1::DsRed (A) and GFP::RAB-11 (B) transgenes in L1 larvae. Images were acquired and displayed with the same settings for comparison. Images are maximum intensity projections. Arrow in (A) indicates a small patch of apical membrane observed at the lateral domain, which occurs with low frequency in ERM-1 T544D mutant animals. (**C,D**) Distribution plots of mean fluorescence intensity of PEPT-1::DsRed (C) and GFP::RAB-11 (D) along the apical-basolateral axis in intestinal cells of L1 larvae. Intensity was normalized to the average peak intensity in wild type animals, and length was normalized to a fixed distance between the basolateral and apical membranes (see methods). Two measurements were done per animal and plotted separately; n≥18 animals. Scale bars: 10 µm.

### Phosphorylation of T544 controls subcellular localization of ERM-1

Interfering with phosphorylation of the C-terminal threonine residue has been reported to affect the subcellular localization of mammalian ezrin and fly Moesin. Replacing the C-terminal threonine with a phosmomimetic aspartic acid or glutamic acid causes relocalization to the entire plasma membrane, while reported effects of alanine substitutions vary from reduced apical enrichment to loss of membrane localization (Coscoy et al., 2002; Karagiosis and Ready, 2004; Roch et al., 2010; Viswanatha et al., 2012; Babich and Di Sole, 2015). We therefore investigated the localization of the phosphorylation mutants in *C. elegans*. To analyze the distribution of wild type and T544 mutant ERM-1 variants we stained embryos and larvae with an antibody directed against *C. elegans* ERM-1 (van Fürden et al., 2004). In the excretory canal, wild type and T544 ERM-1 variants were exclusively detected at the apical/canalicular membrane, suggesting that T544 phosphorylation is not required for subcellular distribution of ERM-1 in this tissue (Fig. S2A). In the intestine, wild type ERM-1 was strongly enriched at the apical plasma membrane throughout embryonic development (Fig. S2B). In contrast, ERM-1[T544A] failed to accumulate at the apical membrane throughout embryogenesis (Fig. S2B). ERM-1[T544D] also failed to accumulate at the apical membrane during early stages of intestine development, but apical enrichment was evident by the 2-fold stage (Fig. S2B). In contrast to wild type ERM-1, which is restricted to the apical membrane soon after intestinal polarization, we observed persistent basolateral localization of ERM-1[T544A] and ERM-1[T544D] throughout embryogenesis. ERM-1 levels in other tissues were difficult to assess due to the quality of antibody staining.

To analyze the distribution of ERM-1 T544 mutants in live animals, we used CRISPR/Cas9 to generate *erm-1[T544A]* and *erm-1[T544D]* alleles carrying a C-terminal GFP fusion. The GFP-tagged T544 mutants showed similar localization patterns as observed by antibody staining. In the intestine, the GFP-tagged T544 mutants showed a delay in apical enrichment (Fig. 6A). However, in contrast to our results using antibody staining, we did observe apical enrichment of ERM-1[T544A]::GFP starting from the 2-fold stage, though at lower levels than ERM-1[T544D]::GFP (Fig. 6A, Fig. S2C). Strikingly, basolateral localization of ERM-1 persisted throughout development in both mutant variants (Fig. 6B,C). Distribution of ERM-1[T544A]::GFP and ERM-1[T544D]::GFP in the excretory canal was analyzed in heterozygous larvae, that form seemingly normal excretory canals. Both ERM-1[T544A]::GFP and ERM-1[T544D]::GFP were detected exclusively at the apical plasma membrane (Fig. 6B). In the germline, ERM-1[T544A]::GFP levels were lower than those of wild-type ERM-1::GFP, but still showed apical enrichment (Fig. S2D,E). In contrast, ERM-1[T544D]::GFP showed increased accumulation at the apical domain (Fig. S2D,E). Thus, C-terminal phosphorylation dynamics modulate ERM-1 distribution in a tissue-specific manner.

**Figure 6.**
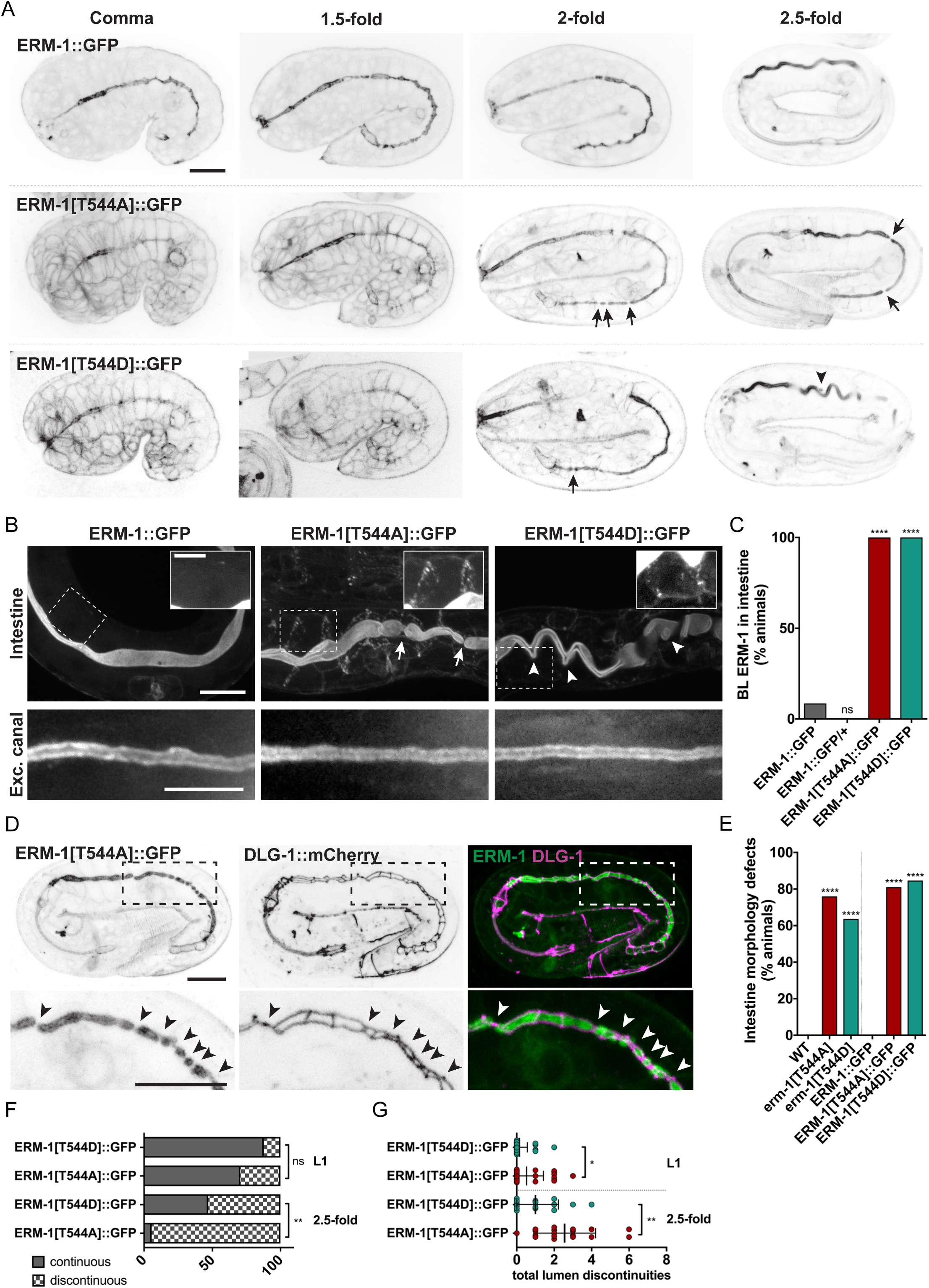
T544 phosphorylation restricts ERM-1 to the apical membrane and drives lumen formation. (**A,B**) Distribution of GFP-tagged ERM-1 variants in different embryonic stages (A) as well as in the intestine and excretory canal of L1 larvae (B). Arrows indicate lumen discontinuities, and arrowheads indicate ectopic ERM-1 localization. Insets are overexposed and enlarged views of the highlighted regions. Excretory canal images are from animals heterozygous for a wild-type *erm-1* allele. (**C**) Frequency of animals with ERM-1 detectable at the basolateral membrane; n≥26 - Fisher’s exact test. Statistical comparisons are with ERM-1::GFP (**D**) Lumen morphology visualized by ERM-1[T544A]::GFP and junction organization visualized by DLG-1::mCherry in a 2.5-fold embryo. Bottom panels are enlarged views of the highlighted regions. Arrowheads indicate lumen discontinuities/junction constrictions. Images are maximum intensity projections. (**E**) Percentage of animals with intestinal morphology defects; n≥26 - Fisher’s exact test. Statistical comparisons are with WT. (**F,G**) Percentage of animals with a continuous or discontinuous lumen (D), and total number of lumen discontinuities per animal in late embryos and L1 larvae (E); n≥17 and n≥34 respectively - Fisher’s exact test (D); Student’s t-test (E). Scale bars: 10 µm; insets in (B) 5 µm.

We also used the *erm-1[T544A]* and *erm-1[T544D]* alleles to examine lumen morphology. In embryogenesis, we observed a delay in separation of opposing apical domains that was more pronounced in ERM-1[T544A]::GFP expressing animals (Fig. 6A,D–G). We also observed constrictions as well as discontinuities along the lumen of *erm-1[T544A]::GFP* L1 larvae, which resolved in later larval stages (Fig. 6A,D–G). The penetrance of intestinal phenotypes was slightly higher than in untagged T544 mutants (Fig. 6E), presumably due to a detrimental influence of the C-terminal GFP tag. These results further demonstrate the importance of ERM-1 phosphorylation in the early stages of intestinal lumenogenesis.

### Mobility of ERM-1 at the membrane is modulated by T544 phosphorylation status

The altered distribution of ERM-1 T544 mutants might be due to changes in the dynamics of ERM-1 association with the membrane. To investigate this, we performed fluorescence recovery after photobleaching (FRAP) experiments using GFP tagged wild-type and T544 mutant strains. In the intestine, wild-type ERM-1::GFP association with the apical membrane was extraordinarily stable, with recovery of bleached areas averaging 15% after 45 minutes (Fig. 7A,B). Analysis of ERM-1[T544A]::GFP and ERM-1[T544D]::GFP forms revealed a more dynamic behavior. In both mutant forms, the fraction and speed of recovery were dramatically increased compared with wild-type ERM-1::GFP, with recoveries averaging 74% for ERM-1[T544A]::GFP and 67% for ERM-1[T544D]::GFP within the same 45 min period (Fig. 7A,B). The slightly slower recovery of ERM-1[T544D]::GFP compared to ERM-1[T544A]::GFP possibly reflects a stronger association with the actin cytoskeleton. Recovery profiles for wild type and T544 mutants were best fitted with a two-component curve, suggesting the presence of two distinct populations (Supplementary Table 1). We did not observe a noticeable difference in recovery at the middle versus the borders of the bleached region (Fig. S3A,B), which suggests that the increased recovery of ERM-1[T544A]::GFP likely results from faster association/dissociation rates, rather than a change in diffusion rate at the membrane.

**Figure 7.**
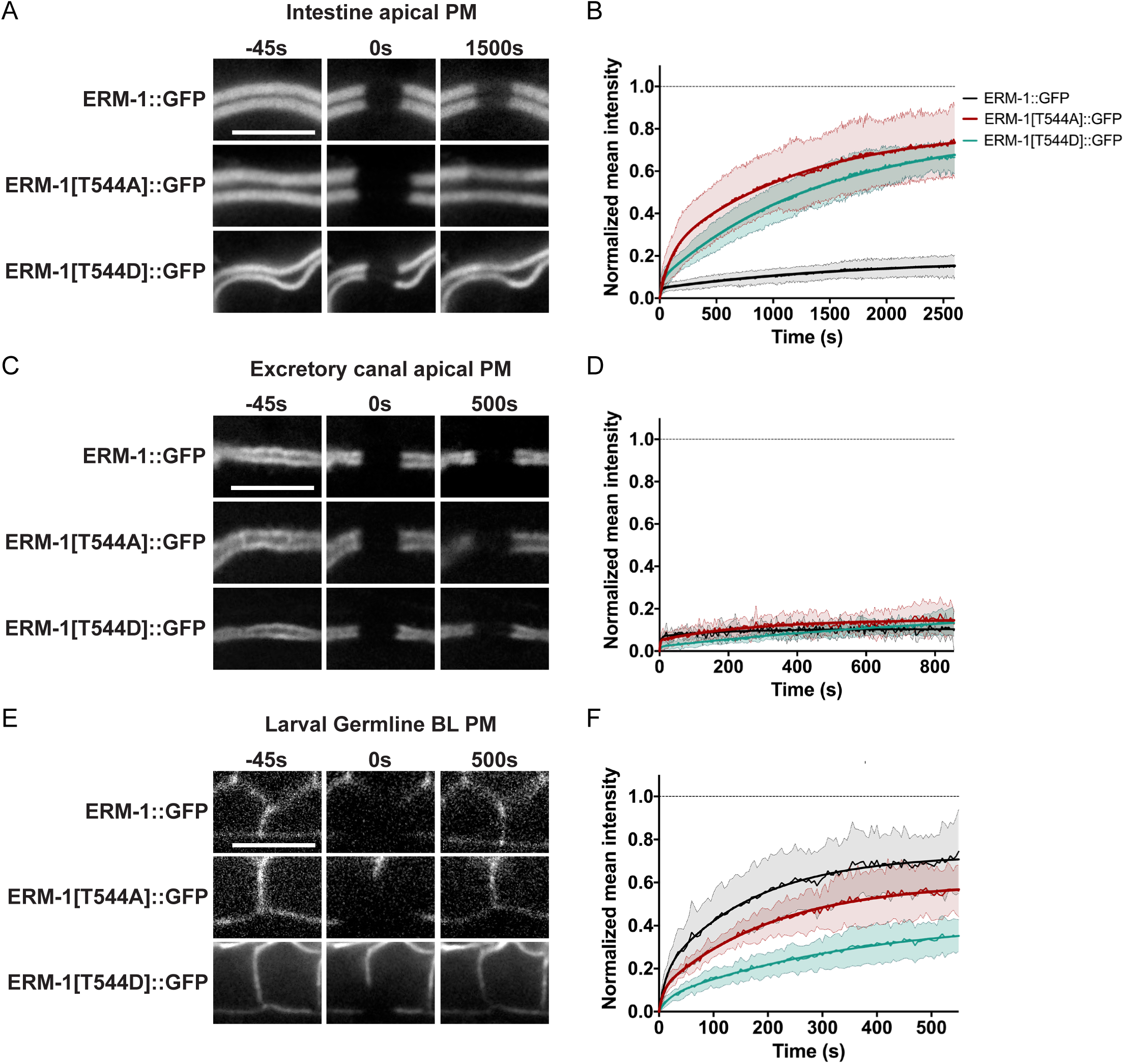
T544 phosphorylation cycling is required for stable ERM-1 localization in the intestine. (**A,B**) Stills from time-lapse movies (A) and FRAP curves (B) of GFP-tagged ERM-1 variants at the apical membrane of L1 larvae intestines; n≥6. In this and subsequent FRAP data thin lines and shading represent the mean ± SD, and thick lines were obtained by curve fitting averaged FRAP data with a double exponential equation. (**C,D**) Stills from time-lapse movies (C) and FRAP curves (D) of GFP-tagged ERM-1 variants at the apical membrane of the excretory canal of L4 larvae heterozygous for wild type *erm-1*; n≥3. (**E,F**) Stills from time-lapse movies (E) and FRAP curves (F) of GFP-tagged ERM-1 variants at the basolateral membrane of the germline of L4 larvae; n≥7. Scale bars: 5 µm.

As C-terminal tagging of ERM-1 disrupts lumen formation in the excretory canal, dynamics of GFP fusions in this tissue were analyzed in heterozygous animals. Recovery rates of wild-type ERM-1::GFP in the excretory canal were similar to the intestine (Fig. 7C,D). However, no changes in recovery rates were detected for T544 mutant forms (Fig. 7C,D). Due to inherent technical challenges in long-term imaging of the excretory canal, we were unable to obtain a satisfactory curve fit for our measurements. We also analyzed the recovery of ERM-1::GFP in the adult germline. Technical challenges prevented us from accurately analyzing the recovery at the apical membrane, thus we focused on the more defined basolateral membrane. ERM-1::GFP was much more dynamic in this tissue, with bleached areas recovering around 70% after 10 minutes (Fig. 7E,F). Surprisingly, both ERM-1[T544A]::GFP and ERM-1[T544D]::GFP had slower recovery profiles (Fig. 7E,F; Supplementary Table 1), suggesting these forms are more stably associated with the basolateral membrane than wild type ERM-1. Our data shows that ERM-1 dynamics are modulated by C-terminal phosphorylation and tissue-specific properties, potentially including different binding partners or differences in lipid membrane composition.

### Phosphorylation of T544 controls apical actin enrichment and dynamics

The C-terminal phosphorylation of ERM proteins has been reported to be required for apical actin enrichment in epithelial cells (Hipfner, 2004; Roch et al., 2010; Abbattiscianni et al., 2016). In *C. elegans*, loss of *erm-1* results in reduced apical actin levels in the intestine and in the excretory canal (Göbel et al., 2004; Bernadskaya et al., 2011; Khan et al., 2013). Furthermore, overexpression of ERM-1 in the excretory canal leads to excessive accumulation of apical actin coating the lumen (Khan et al., 2013). The apical actin network of tubular epithelia in *C. elegans* is mostly composed of the specialized actin ACT-5 (MacQueen et al., 2005). To analyze ACT-5 in the canal, we generated a transgenic line expressing mCherry::ACT-5 from the *sulp-4* promoter. Consistent with our results using VHA-5::GFP, the excretory canals showed variable, but fully penetrant, morphological defects in both ERM-1 T544 mutants (Fig. 8A). In canal regions with a widened lumen, apical ACT-5 coating was sparser and circumferential bundles were visible. The apical to cytoplasm ratio of mCherry::ACT-5 was decreased in both *erm-1[T544A]* and *erm-1[T544D]* mutants, relative to wild type (Fig. 8B). Thus, in the excretory canal, cycling of T544 phosphorylation appears to be important for ERM-1 to recruit actin to the apical membrane.

**Figure 8.**
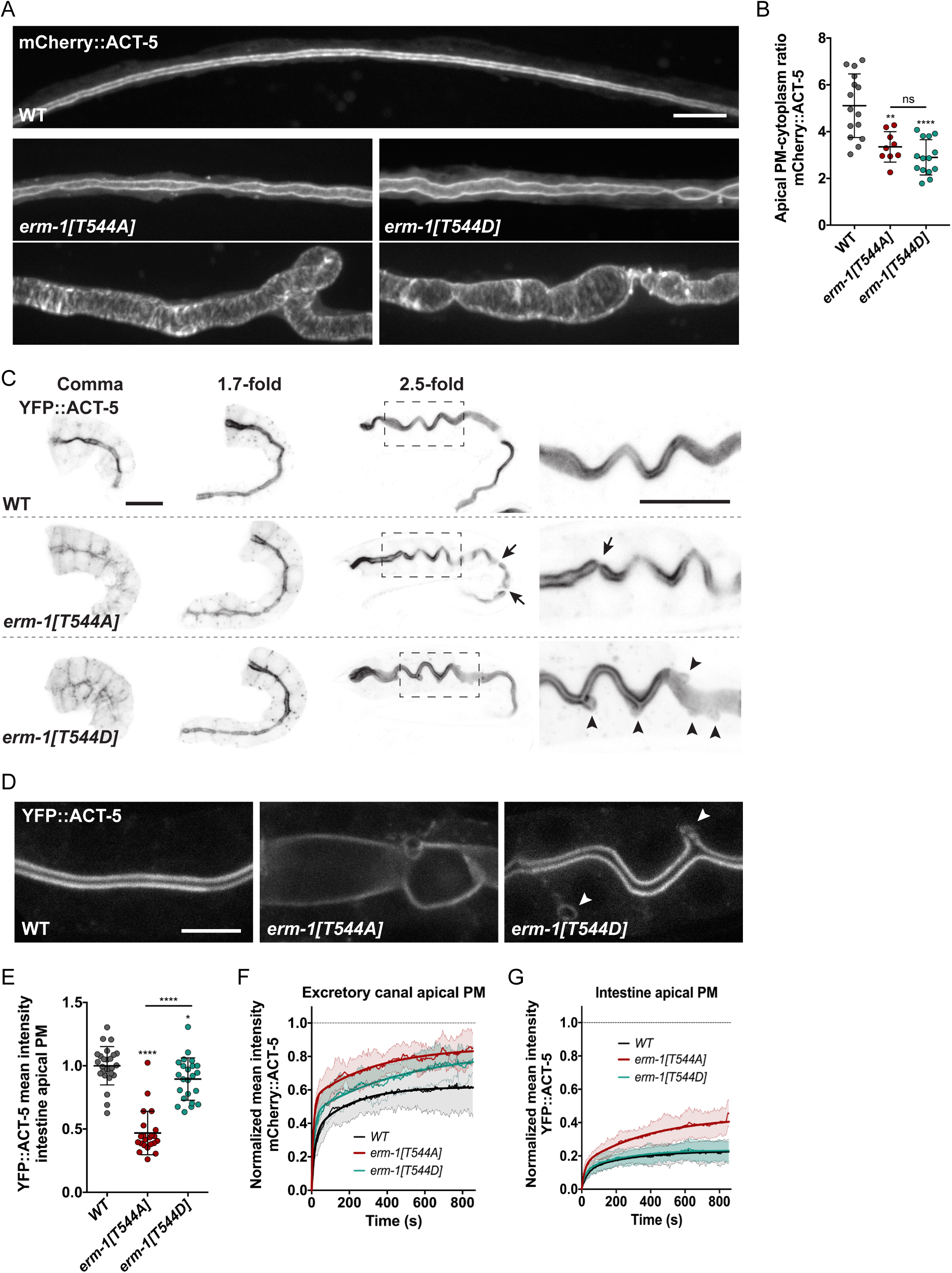
Phosphorylation of T544 controls apical actin recruitment and dynamics. (**A**) Lumen morphology and distribution of mCherry::ACT-5 in the excretory canal of L4 larvae. Bottom panels are examples of severe phenotypes in *erm-1[T544A]* and *erm-1[T544D]* mutants. (**B**) Apical/cytoplasmic mCherry::ACT-5 ratio of mean intensity in the excretory canal of L4 larvae. Ratio was based on the average of three measurements at the apical membrane and cytoplasm, and each data point represents one animal; n≥9. Unless indicated otherwise by a connecting line, statistical comparisons are with WT. (**C,D**) Intestinal distribution of YFP::ACT-5. Panels to the right in (C) are enlarged views of the highlighted regions. Arrows point to discontinuities along the cortical actin cytoskeleton, and arrowheads point to ectopic expansions of the cortical actin network. Scale bars: 10 µm. (**E**) Plots of intestinal YFP::ACT-5 mean intensity at the apical membrane. Each symbol represents a single animal; n≥22. Unless indicated otherwise by a connecting line, statistical comparisons are with WT. (**F,G**) FRAP curves of apical mCherry::ACT-5 in the excretory canal (G; n≥7), and apical YFP::ACT-5 in the intestine (H, n≥8), of L1 larvae.

To analyze ACT-5 distribution in intestinal cells we used an integrated YFP::ACT-5 transgene (Bossinger et al., 2004). In intestines of wild type embryos, we detected a strong enrichment of ACT-5 at the apical domain of intestinal cells soon after polarization (Fig. 8C). In contrast, we failed to detect significant apical enrichment of YFP::ACT-5 in comma stage embryos expressing either ERM-1[T544A] or ERM-1[T544D] (Fig. 8C), and YFP::ACT-5 was clearly detectable in the cytoplasm and along the basolateral membrane. Similar to the distribution of ERM-1 T544 mutants during embryogenesis, we did observe apical enrichment of YFP::ACT-5 in *erm-1[T544A]* and *erm-1[T544D]* animals at later embryonic stages, which was maintained throughout development (Fig. 8C,D; Fig. S3C). Nevertheless, quantification of YFP::ACT-5 distribution in *erm-1[T544A]* L1 larvae showed a severe reduction in apical levels (Fig. 8D,E). In contrast, ACT-5 levels in animals expressing ERM-1[T544D] were unaffected (Fig. 8D,E).

We next investigated ACT-5 dynamics by FRAP in L1 animals. The effects on ACT-5 recovery by *erm-1[T544A]* and *erm-1[T544D]* mutations were consistent with the effects we observed on ACT-5 levels. In the excretory canal, recovery of mCherry::ACT-5 was slightly faster in both mutants than in *erm-1* wild type animals, with the fastest recovery observed in the *erm-1[T544A]* background (Fig. 8F). In the intestine, recovery of YFP::ACT-5 in wild type animals was slow (∼23% in 15 minutes) and similar to that of ERM-1::GFP. Recovery of apical YFP::ACT-5 in the intestine of *erm-1[T544A]* mutants was faster than in wild type animals (∼41% in 15 minutes), whereas no difference was seen in *erm-1[T544D]* (Fig. 8G). Although we did not observe any defects in actin levels from the L2 stage onward (Fig S3C), intestinal ACT-5 still recovered faster in *erm-1[T544A]* L4 mutants compared to wild type (Fig. S3D).

Collectively, our results suggest that T544 phosphorylation is important for cortical actin recruitment by ERM-1 in tubular epithelia. In the excretory canal, accumulation of cortical actin requires T544 phosphorylation cycling, while in the intestine this process is mostly dependent on the phosphorylated form of ERM-1. These observations also indicate that phosphorylated ERM-1 forms are better able to interact with actin, which is consistent with biochemical analysis of ERM phosphorylation mutants *in vitro* and in mammalian cells (Gautreau et al., 2000; Bosk et al., 2011; Braunger et al., 2014). Finally, the similar delay in apical accumulation of ERM-1 and ACT-5 in the intestine of T544 mutants suggest that presence of ERM-1 at the apical membrane is the major factor in ACT-5 recruitment.

## Discussion

Activation of ERM proteins at spatially controlled regions of the cortex drives morphological specialization events required for the function of numerous cell-types across animal species. Here, we characterized the contribution of the major conserved regulatory sites important for the activity of ERM proteins *in vivo* using *C. elegans*. The use of CRISPR/Cas9 allowed us to analyze the effects of PIP_2_-binding and C-terminal phosphorylation mutants in the absence of wild type product, without addition of any tags, and at endogenous expression levels. This is especially relevant considering that ERM proteins are known to form stable dimers and oligomers, are affected by fluorescent protein tags, and often have dosage-dependent effects (Berryman, 1995; Gautreau et al., 2000; Cao et al., 2005; Chambers and Bretscher, 2005; Khan et al., 2013). In agreement with studies in *Drosophila* and mammalian cultured cells (Yonemura et al., 2002; Fievet et al., 2004; Hao et al., 2009; Roch et al., 2010), we show that PIP_2_-binding is essential for ERM-1 activity. In contrast, our ‘clean’ phosphorylation mutants demonstrate that phosphorylation is not essential for ERM activity in *C. elegans*, despite contributing to multiple aspects of ERM-1 function, often in a tissue-specific manner.

We analyzed the contribution of PIP_2_-binding for ERM-1 activity by mutating four lysine residues within the FERM domain, whose importance for PIP_2_-binding has been well established (Hirao, 1996; Barret et al., 2000; Fievet et al., 2004; Hao et al., 2009; Li et al., 2007; Ben-Aissa et al., 2012; Jayasundar et al., 2012; Babich and Di Sole, 2015; Roch et al., 2010). The similarity in phenotypes caused by the *erm-1[4KN]* mutation and the putative *erm-1(tm667*) null allele (Göbel et al., 2004) demonstrates the essentiality of PIP_2_ binding for ERM-1 activity. Effects on protein localization were also largely consistent with an essential role for PIP_2_ binding in localizing ERM-1 to the plasma membrane. Unexpectedly, however, in heterozygous *erm-1[4KN]::GFP/+* animals and in first generation homozygous animals we detected substantial amounts of ERM-1 at the plasma membrane of intestinal cells and the excretory canal. To explain this observation, we considered the possibility that ERM-1 localizes to the plasma membrane as a dimer. However, first generation *erm-1[4KN]::GFP* homozygous animals contain very low levels of wild-type ERM-1. Thus, formation of ERM-1 dimers does not appear to be able to account for the significant plasma membrane localization we observed. An alternative model is the formation of higher order ERM-1 complexes, in which the presence of only a few wild-type ERM-1 molecules would be sufficient to mediate membrane localization. Finally, wild-type ERM-1 may enact apical membrane modifications that are sufficient to recruit ERM-1[4KN]. In agreement with a wild type recruitment model, co-fractionation assays done in the presence of endogenous ezrin still detected a substantial amount of ezrin-4KN in the membrane fraction (Barret et al., 2000; Fievet et al., 2004; Babich and Di Sole, 2015).

To investigate the contribution of phosphorylation on the conserved C-terminal threonine residue (T544 in ERM-1) to ERM activity, we mutated the endogenous *erm-1* gene to express either a non-phosphorylatable T544A variant or a phosphomimetic T544D variant. Unexpectedly, our results show that C-terminal phosphorylation is not essential for ERM-1 functioning in *C. elegans*, despite contributing to the regulation of ERM-1 in different tissues. Perhaps most strikingly, both ERM-1 T544A and T544D animals develop intestines with a normal appearance, and only a mildly reduced length of microvilli. These results contrast with multiple studies in mammalian cell culture and *Drosophila*, which reported that non-phosphorylatable and phosphomimetic ERM variants cannot substitute for the activity of wild-type ERM proteins (Viswanatha et al., 2012; Parameswaran et al., 2011; Polesello et al., 2002; Karagiosis and Ready, 2004). However, in support of our observations, one study in *Drosophila* showed that expression of a Moesin-T559A transgene (but not Moesin-T559D) could significantly rescue viability of a strong *moesin* allele (Roch et al., 2010), and two others demonstrated rescuing activity for Moesin-T559D – though not for Moesin-T559A (Hipfner, 2004; Speck et al., 2003).

The reduced microvilli length in T544A animals may be due to a less efficient recruitment of actin, as L1 stage larvae display severely reduced levels of apical actin. However, T544D L1 larvae display near normal levels of apical actin, yet develop intestines with shorter microvilli than T544A mutant animals. This indicates that actin recruitment is not the only function of ERM-1 in supporting microvilli development. The severely reduced levels of PEPT-1 in both ERM-1 phosphorylation mutants are consistent with an essential role for ERM-1 in specifying the composition of the apical domain, which may well include factors that affect microvilli growth. Although not directly addressed in our study, it is likely that non-phosphorylatable ERM-1[T544A] can still interact (inefficiently) with actin, as the linker function is at the core of ERM activity and we observe significant actin recruitment in multiple tissues. Indeed, it was previously shown that non-phosphorylatable forms can still interact with actin *in vitro* (Matsui et al., 1998; Bosk et al., 2011; Braunger et al., 2014). Finally, basolateral localization of ERM-1, as caused by both phosphorylation mutants, might be expected to result in conversion of basolateral to apical domain identity. Although we noticed an occasional lateral ‘bubble’ of apical membrane in T544D mutant animals (arrow in Fig. 5A), we did not observe widespread changes to basolateral identity. However, basolateral levels of ERM-1 are very low, and the mutation of T544 may reduce the ability to specify apical identity. Hence, whether *C. elegans* ERM-1 is sufficient to induce apical domain identity cannot be concluded from our experiments.

The defects induced by expression of ERM-1[T544D] or ERM-1[T544A] were highly similar. This includes the lumen formation defects in both the excretory canal and early intestine, microvilli formation in intestinal cells, and the defects in ERM-1 distribution and mobility. These results are consistent with a model in which cycling between phosphorylated and non-phosphorylated states is crucial to controlling the activity of ERM-1, as has been proposed to account for ERM protein activity in activation of B and T cells, in secretion of gastric parietal cells, and microvilli formation in epithelial cells (Zhu et al., 2007; Parameswaran et al., 2011; Viswanatha et al., 2012). This interpretation does rely on the assumption that phosphomimetic mutations mimic the phosphorylated state. This assumption is supported by extensive characterization of phosphomimetic mutants in other systems (Huang et al., 1999; Matsui et al., 1998; Nakamura et al., 1999; Oshiro et al., 1998; Simons et al., 1998; Gautreau et al., 2000; Polesello et al., 2002; Speck et al., 2003; Chambers and Bretscher, 2005; Charras et al., 2006; Carreno et al., 2008; Kunda et al., 2008; Bosk et al., 2011), and by our finding that the phosphomimetic ERM-1[T544D] form is recognized by the pERM antibody. Our data therefore strongly advocates that dynamic turn-over of C-terminal phosphorylation is a key mechanism regulating ERM activity.

Our analysis of ERM-1::GFP protein dynamics using FRAP revealed an unexpected stability for ERM-1 in the intestine and excretory canal. Previous FRAP studies in microvilli of LLC-PK1 cells and in blebbing M2 melanoma cells indicate the presence of three pools of ezrin with recovery half-times in the range of seconds to 1–2 minutes (Coscoy et al., 2002; Fritzsche et al., 2014). Curve fitting with exponential functions also supports the presence of at least two pools of ERM-1 in *C. elegans*. However, we observed little fluorescence recovery even after 45 min in either tissue, and recovery half-times in the intestine for the two ERM-1 pools were 8.3 and 1276 seconds, respectively. Slow recovery of ERM-1 is consistent with a recent study examining the interaction of actin binding proteins with phosphoinositide-rich membranes, which found that the ezrin FERM domain binds membranes with very high affinity and slow dissociation dynamics (Senju et al., 2017). In contrast to the intestine and excretory canal, ERM-1::GFP levels at the basolateral domain of the gonad recovered to ∼70% within 10 minutes, with recovery half-times of 8.4 and 117 s respectively, within the range of previous observations in mammalian cells. Interestingly, T544A and T544D substitutions greatly increased the mobility of ERM-1::GFP in the intestine. How can phosphorylation cycling result in a more stable apical localization of ERM-1? While it is difficult to speculate, ERM-1 may need to cycle between different conformations in order to cope with a changing microenvironment, that presumably involves actin treadmilling, and local changes in the concentration or availability of PIP_2_ and various protein binding partners. In contrast to the intestine, ERM-1 stability in the excretory canal was unaffected by the phosphorylation mutants, perhaps reflecting a difference in the dynamics of apical membrane components between tissues. Regardless of the underlying reason, our observations demonstrate that ERM-1 turnover dynamics can vary greatly dependent on biological context, and are subject to tissue-specific regulation, which includes but is not limited to T544 phosphorylation.

The tissue-specific effects of T544 mutations extended to tissue formation and localization of ERM-1. In the intestine, both *erm-1[T544A]* and *erm-1[T544D]* animals ultimately form a continuous intestinal tube containing microvilli, in contrast to *erm-1* loss of function. However, both T544 mutations led to dramatically increased mobility and ubiquitous cortical distribution of ERM-1. In early stages of intestinal development, we observed a delay in apical recruitment of ERM-1, a delay in apical enrichment of actin, and a delay in lumen formation resembling previous descriptions of *erm-1* mutant alleles and RNAi (van Fürden et al., 2004; Göbel et al., 2004). Based on the similarity in the timing of these events we hypothesize that phosphorylation cycling is crucial for early stages of intestinal lumen formation by assuring the timely apical enrichment and basolateral exclusion of ERM-1. Support for phosphorylation cycling-dependent local stabilization at the apical plasma membrane comes from a study by Viswanatha and colleagues, which showed that localized kinase activity at the apical membrane of microvilli-containing mammalian cells is able to restrict ezrin to the apical plasma membrane (Viswanatha et al., 2012). Similar polarized kinase activity was shown to restrict Moesin activity to the apical cortex of tracheal cells in Drosophila (Ukken et al., 2014). Phosphorylation-dependent apical recruitment may represent a general mechanism of localized ERM enrichment and activation as different studies have shown that altered C-terminal phosphorylation perturbs the polarized distribution of ERM proteins (Coscoy et al., 2002; Polesello et al., 2002; Dard et al., 2004; Fievet et al., 2004; Karagiosis and Ready, 2004; Viswanatha et al., 2012).

In contrast to the intestine, lumen formation and outgrowth of the excretory canal were severely affected in both *erm-1[T544A]* and *erm-1[T544D]* mutant animals throughout the lifespan of the animal. These results confirm that excretory canal formation is strongly dependent on ERM-1 activity. Indeed, ERM-1 has been shown to control canal lumen formation in a dose-dependent manner, whereas no effect has been observed for ERM-1 overexpression in the intestine (Göbel et al., 2004; Khan et al., 2013). Surprisingly, however, neither localization nor mobility of ERM-1 in the canal were affected by altered T544 regulation. ERM proteins are known to interact with and regulate the function of numerous proteins important for the formation of biological tubes, including trafficking components, transmembrane channels or pumps, polarity determinants, and junction proteins (Médina et al., 2002; Deretic et al., 2004; Pilot et al., 2006; Zhu et al., 2007; Chirivino et al., 2011; Kvalvaag et al., 2013; Khan et al., 2013; Bryant et al., 2014). Moreover, several interaction partners have been shown to selectively interact with either non-phosphorylated or phosphorylated forms of ezrin (Viswanatha et al., 2013). It is possible therefore that a difference in the subset of ERM-1 interaction partners involved in each tissue explains why effects on ERM-1 localization and mobility do not strictly correlate with the phenotypic defects at the tissue level. The absence of changes in ERM-1 protein behavior in the canal further indicate that C-terminal threonine phosphorylation is not the only mechanism that controls ERM-1 localization and stability. Binding to specific interaction partners may contribute to ERM-1 localization, but alternative mechanisms of regulation are also possible. For example, a recent study found that in breast epithelial cells, ezrin membrane association is regulated by acetylation of ezrin (Song et al., 2019). Overall, these results demonstrate that C-terminal phosphorylation is a versatile regulatory modification that modulates ERM-1 function in a context-dependent manner and underscore the importance of studying ERM regulation in different biological scenarios.

## Materials and Methods

### *C. elegans* strains and culture conditions

*C. elegans* strains were cultured under standard conditions (Brenner, 1974). Only hermaphrodites were used and all experiments were performed with animals grown at 20 °C on Nematode Growth Medium (NGM) agar plates. Supplementary Table 2 contains a list of all the strains used.

### CRISPR/Cas9 genome engineering

Endogenous eGFP protein fusions and point mutations were generated by homology-directed repair of CRISPR/Cas9-induced DNA double-strand breaks (DSBs). *erm-1[T544A]* and eGFP proteins fusions were generated in an N2 background, with the exception of *erm-1[4KN]::GFP*, in which *erm-1[KN]* was used as the starting strain; remaining point mutations were generated in a *pha-1(e2123ts)* background. In all cases two sgRNA plasmids targeting each locus were used. sgRNA plasmids were generated by ligation of annealed oligo pairs into the *pU6::sgRNA* expression vectors pMB70 (Addgene #47942) or pJJR50 (Addgene #75026) as previously described (Waaijers et al., 2013, 2016). To introduce point mutations synthesized single-stranded oligodeoxynucleotides (ssODNs) with 33-45 bp homology arms were used as a repair template, and integration events were selected using either *dpy-10* (Arribere et al., 2014) or *pha-1* (Ward, 2015) co-CRISPR approaches. eGFP knock-ins were introduced using a plasmid-based repair template with 450-600 bp homology arms and containing a self-excising cassette (SEC) for selection, as previously described (Dickinson et al., 2015). To introduce eGFP we created a custom SEC vector, pJJR82 (Addgene #75027), by replacing a fragment of pDD282 (Addgene #66823) comprising the GFP sequence with a similar fragment comprising a codon optimized and synthetic intron-containing eGFP sequence using the flanking Bsu36I and BglII restriction sites. In all cases, correct genome editing was confirmed by Sanger sequencing (Macrogen Europe) of PCR amplicons encompassing the edited genomic region.

The maternal-effect lethal *erm-1[KN]* and *erm-1[KN]::GFP* alleles were balanced with *dpy-5(e61)*; *unc-29(e403)* I (DR102 strain). In all cases edited strains were backcrossed twice with N2 to eliminate any non-linked unspecific editing events, and one additional round of backcrossing was done for strains generated in a *pha-1(e2123ts)* background. The sequences of all oligonucleotides used (synthesized by Integrated DNA technologies) are listed in Supplementary Table 3.

### Microscopy

Imaging of *C. elegans* was done by mounting embryos or larvae on a 5% agarose pad in a 10 mM Tetramisole solution in M9 buffer (0.22 M KH_2_PO_4_, 0.42 M Na_2_HPO_4_, 0.85 M NaCl, 0.001 M MgSO_4_) to induce paralysis. Spinning disk confocal imaging was performed using a Nikon Ti-U microscope equipped with a Yokogawa CSU-X1 spinning disk and an Andor iXon+ EMCCD camera, using a 60x-1.4 NA objective. Time-lapse imaging for FRAP experiments was performed on a Nikon Eclipse-Ti microscope equipped with a Yokogawa CSU-X1-A1 spinning disk and a Photometrics Evolve 512 EMCCD camera, using a 100x 1.4 NA objective. Targeted photobleaching was done using an ILas system (Roper Scientific France/ PICT-IBiSA, Institut Curie). Two epifluorescence microscopy set-ups were used. DIC imaging was done in an upright Zeiss Axioplan2 microscope using a 63x-1.4 NA objective, while imaging to quantify excretory canal outgrowth was done in an upright Zeiss Axio Imager 2 microscope using a 20x-0.5 NA objective. Microscopy data was acquired using MetaMorph Microscopy Automation & Image Analysis Software (spinning Disk), Zeiss AxioVision (DIC), and Zeiss Zen (epifluorescence). All stacks along the z-axis were obtained at 0,25 μm intervals, and all images were analyzed and processed using ImageJ and Adobe Photoshop.

### Quantitative image analysis

Quantitative analysis of spinning disk images was done in ImageJ. In all quantifications, mean background intensity was quantified by drawing a circular region of 50 px diameter in areas within the field-of-view that did not contain any animals, and values were normalized using the mean intensity of eGFP, YFP, or mCherry at the apical membrane of the corresponding tissue in control animals. In the intestine, all measurements were done in cells forming int2 through int6, and fluorescence intensity at the apical membrane was quantified in regions where opposing apical membranes could be clearly seen as two lines. To quantify fluorescence intensity of ERM-1[4KN]::GFP in intestinal cells and mCherry::ACT-5 in the excretory canal, measurements were performed in maximum intensity projections of three consecutive z-slices showing the highest intensity at the apical membrane. Peak intensity at the apical membrane was calculated by averaging the peak values of intensity profiles from multiple 40 px-wide line-scans perpendicular to the membrane per animal. Mean cytoplasmic intensity was obtained by averaging the mean intensity values of multiple elliptical regions within the cytoplasm. Each measurement was corrected for background noise and normalized as described above. Averaged apical and cytoplasmic intensity values were used to calculate the apical to cytoplasmic intensity ratio per animal. Distribution plots of fluorescence intensity of eGFP-tagged ERM-1 variants in the larval germline as well as PEPT-1::DsRed and GFP::RAB-11 in the intestine, were obtained using a similar method. For eGFP-tagged ERM-1 variants in germline, three measurements were done per animal in single frames where both apical and basal membranes were clearly visible. For animals expressing both PEPT-1::DsRed and GFP::RAB-11 in the intestine, two measurements were done per animal in maximum intensity projections of the three consecutive z-slices with the highest intensity of PEPT-1::DsRed at the apical membrane. Measurements from single animals were not averaged in either case. Intensity distribution profiles were obtained by tracing line-scans (15 px-wide for the germline and 40 px-wide for the intestine) encompassing the entire germ cell compartments or intestinal cells along the apical-basal axis. Intensity profiles were manually trimmed to exclude values outside the cells/compartments of interest, and each value was corrected for background noise and normalized as described above. To directly compare and plot intensity profiles despite differences in the distance between basal and apical membranes, a custom R script was made to linearly interpolate intensity values in the y axis to a fixed distance along the x axis for each intensity profile defined by the average apical-basal distance in control animals (script is available upon request). For the germline, apical-basal intensity ratios were calculated using the peak intensity values at the apical and basal membranes per intensity profile. To quantify fluorescence intensity of eGFP-tagged ERM-1 variants and YFP::ACT-5 in intestinal cells, a free-hand region was drawn either surrounding the apical membrane or in the cytoplasm, and mean intensity values were extracted for all z-slices in which apical membrane was visible. The background was subtracted per frame and each value was normalized as described above. Mean intensity at the apical membrane and cytoplasm were calculated by averaging measurements through the z axis of two intestinal cells per animal. Averaged intensity values were used to calculate apical-cytoplasm ratio per animal.

### Brood size and lethality

Starting at the L4 stage, individual P0 animals were cultured at 20 °C and transferred to a fresh plate every 24 h for 6 days. Hatched and unhatched progeny were scored 24 h after removal of the P0, and larval lethality was scored 48 h after removal of the P0.

### Relative excretory canal outgrowth

To quantify relative canal outgrowth in the excretory canal cell, F1 progeny of L4 animals expressing the VHA-5::GFP transgene grown in standard or RNAi culture plates was scored at the L4 stage. The distance between the cell body and either anterior or posterior distal body tips was determined by tracing a segmented line along the center of the animal. Length of each individual canal was measured with a segmented line from the anterior-posterior bifurcation points close to the cell body until the canal tip. Relative outgrowth was calculated as the fraction of canal length over the distance between the cell body and distal tips. Severe outgrowth defects were defined as canals that extend 35 % or less of the distance between the excretory canal cell body and either anterior or posterior tips. Frequency was calculated by the sum of both anterior and posterior canals with severe defects over total canals quantified per genotype.

### FRAP experiments and analysis

For FRAP assays, laser power was adjusted in each experiment to avoid complete photobleaching of the selected area, as the time scale of experiments prevented assessment of photo-induced damage. Photobleaching was performed on a circular region with a diameter of 30 or 40 px at the cortex, and recovery was followed at 5 s intervals for 15-45 min depending on the tissue. Time-lapse movies were analyzed in ImageJ. The size of the area for FRAP analysis was defined by the full-width at half maximum of an intensity plot across the bleached region in the first post-bleach frame. For each time-lapse frame, the mean intensity value within the bleached region was determined, and the background, defined as the mean intensity of a non-bleached region outside the animal, was subtracted. The mean intensities within the bleached region were corrected for acquisition photobleaching per frame using the background-subtracted mean intensity of a similar non-bleached region at the cortex, which was normalized to the corresponding pre-bleach mean intensity. FRAP recovery was calculated as the change in corrected intensity values within the bleach region from the first frame after bleach (set to normalized to the mean intensity of the 10 frames before bleach. Curve fitting was done on averaged recovery data per sample using the non-linear regression analysis on GraphPad. One and two-phase association were tested and in all cases data were best fitted with a two-phase curve. Intensity distribution plots were obtained by performing a 3 px wide line-scan perpendicular to the apical membrane.

### Immunohistochemistry

Antibody stainings were performed in mixed-stage (anti-pERM) or synchronous (anti-ERM-populations. Synchronized animals were obtained from gravid adult animals by bleaching and allowed to develop in M9 for 4-6 h for embryonic stages, or in standard culture plates for 10 h for early larval stages. Mixed-stage or synchronous larval populations were collected from plates and were washed 3 times in M9 (for stainings of post-embryonic stages, samples were incubated for 30 minute with gentle shaking before the last wash), washed once in water, and transferred to a poly-L-Lysine coated slide. Samples were permeabilized by freeze-cracking and fixed at −20 °C with methanol for 5 min and acetone for 10 min for staining with the ERM- 1 antibody, or with P buffer (3.7 % formaldehyde, 75 % methanol, 250 µM EDTA, 50 mM NaF) for 15 min and methanol for 5 min for the pERM antibody. Samples were rehydrated in an ethanol series (90 %, 60 %, and 30 %, for 10 min each at −20 °C), rinsed three times in PBST (1.35 M NaCl, 27 mM KCl, 100 mM Na2HPO4, 18 mM KH_2_PO_4_, 0.05 % Tween-20), and blocked for 1 h with 1 % BSA and 10 % serum in PBST at room temperature (RT). Samples were incubated with primary antibodies in blocking solution overnight at 4 °C, washed four times for 15 minutes in PBST, and secondary antibodies in blocking solution were incubated for 2 h at RT. Finally, samples were washed three times in PBST, and once in PBS for 10 min each and mounted with Prolong Gold Antifade (Thermofisher). The following antibodies and dilutions were used: MH27 mouse monoclonal (Developmental Studies Hybridoma Bank) 1:20, ERM-1 rabbit polyclonal (gift from O. Bossinger) 1:100, pERM rabbit polyclonal (Cell Signaling Technologies) 1:100, Alexa-Fluor 488 goat anti-rabbit, and Alexa-Fluor 568 goat anti-mouse (Thermofisher) 1:500.

### Feeding RNAi

The *aqp-8* RNAi clone was obtained from the genome wide Vidal full-length HT115 RNAi feeding library derived from the ORFeome 3.1 collection (Rual et al., 2004). An HT115 bacterial clone expressing the L4440 vector lacking an insert was used as a control. For feeding RNAi experiments, bacteria were pre-cultured in 2 ml Lysogeny Broth (LB) supplemented with 100 µg/ml ampicilin (Amp) and 2.5 µg/ml tetracyclin (Tet) at 37 °C in an incubator rotating at 200 rpm for 6-8 h, and then transferred to new tubes with a total volume of 10 mL for overnight culturing. To induce production of dsRNA, cultures were incubated for 90 min in the presence of 1 mM Isopropyl β-D-1-thiogalactopyranoside (IPTG). Bacterial cultures were pelleted by centrifugation at 4000 g for 15 min and concentrated 5x. NGM agar plates supplemented with 100 μg/ml Amp and 1 mM IPTG were seeded with 250 μl of bacterial suspension, and kept at room temperature (RT) for 48 h in the dark. Six to eight L4 hermaphrodites per strain were transferred to individual NGM-RNAi plates against target genes and phenotypes were analyzed in the F1 generation.

### Transmission Electron Microscopy

For transmission electron microscopy (TEM), L4 animals were fixed by high-pressure freezing with the EMPACT-2 system (Leica Microsystems). Freeze-substitution (FS) was done in anhydrous acetone containing 1 % OsO_4_, 0.5 % glutaraldehyde and 0.25 % uranyl acetate for 60 h in a FS system (AFS-2, Leica Microsystems). Larvae were embedded in an Epon-Araldite mix (hard formula, EMS). Adhesive frames were used (Gene Frame 65 μl, ThermoFisher) for flat-embedding, as previously described (Kolotuev et al., 2012), to facilitate anterior-posterior orientation and sectioning. Ultrathin sections were cut on an ultramicrotome (UC7; Leica Microsystems) and collected on formvar-coated slot grids (FCF2010-CU, EMS). Each larvae was sectioned in five different places with ≥10 μm between each grid to ensure that different cells were observed. Each grid contained at least ten consecutive sections of 70 nm. TEM grids were observed using a JEM-1400 transmission electron microscope (JEOL) operated at 120 kV, equipped with a Gatan Orius SC200 camera (Gatan) and piloted by the Digital Micrograph program. Microvilli length was quantified using Fiji on TEM pictures of at least five sections per worm.

### Excretory canal-specific mCherry::ACT-5 reporter

The *Psulp-4::mCherry::ACT-5* construct was cloned into the pBSK vector using Gibson Assembly (Gibson et al., 2009). A fragment containing 2.3 kb immediately upstream of the *sulp-4* coding sequence, and a fragment of 1.7 kb containing the entire genomic sequence of *act-5* and 215 bp of the 3’ UTR, were amplified from *C. elegans* genomic DNA. The codon-optimized mCherry sequence with synthetic introns and a C-terminal linker was amplified from pJJR83 (Addgene #75028). Correct amplification and assembly was confirmed by Sanger sequencing. Primers used are listed in Supplementary table 3. Several stable transgenic lines were generated by microinjection of N2 young adult animals with 5 ng/µl *Psulp-4::mCherry::ACT-5* and 10 ng/µl *Plin-48::GFP* as a co-injection marker, which did not affect excretory canal development, and one was selected for further analysis.

### Statistical analysis

Statistical tests used and sample sizes are indicated in the figure legends and in the methods section. No statistical method was used to pre-determine sample sizes. No samples or animals were excluded from analysis. The experiments were not randomized, and the investigators were not blinded to allocation during experiments and outcome assessment.

## Acknowledgments

We thank J. Kerver-Stumpfova for technical assistance, H. R. Pires, J. Cravo and S. van den Heuvel for providing the *dlg-1::mCherry; hmr-1::eGFP* knock-in strain, O. Bossinger for providing the ERM-1 antibody and the BJ49 strain, M. Labouesse for sharing the ML846 strain, and M. Zerial for sharing the MZE1 strain. We acknowledge Wormbase, and the Biology Imaging Center Faculty of Sciences, Department of Biology, Utrecht University. Some strains were provided by the Caenorhabditis Genetics Center, which is funded by NIH Office of Research Infrastructure Programs (P40 OD010440). The MH27 monoclonal antibody developed by R. Francis and R. H. Waterston was obtained from the Developmental Studies Hybridoma Bank, created by the NICHD of the NIH and maintained at The University of Iowa. We thank S. van den Heuvel, M. Harterink and members of the S. van den Heuvel, M. Boxem, and R. Korswagen groups for helpful discussions. This work was supported by the Netherlands Organization for Scientific Research (NWO)-CW ECHO 711.014.005 and NWO-VICI 016.VICI.170.165 grants to M. Boxem, and by funding from the Ligue contre le Cancer Grand-Ouest/22-29, the Centre National de la Reserche Scientifique, and Université Rennes to G. Michaux.

## Author Contributions

João J. Ramalho contributed to conceptualization, formal analysis, investigation, methodology, visualization, and writing of the manuscript. Ophélie Nicolle contributed to formal analysis, investigation, methodology and visualization. Grégoire Michaux contributed to funding acquisition, supervision, review and editing. Mike Boxem contributed to conceptualization, funding acquisition, project administration, supervision, visualization, and writing of the manuscript.

