## Supplementary Tables and Figures for "*In vivo* control of the ezrin/radixin/moesin protein ERM-1 in *C. elegans*"

### Supplementary Material

João J. Ramalho[1,3], Ophélie Nicolle[2], Grégoire Michaux[2], and Mike Boxem[1]

1 – Division of Developmental Biology, Institute of Biodynamics and Biocomplexity, Department of Biology, Faculty of Science, Utrecht University, Padualaan 8, 3584 CH, Utrecht, The Netherlands

2 – Univ Rennes, CNRS, IGDR (Institut de Génétique et de Développement de Rennes), UMR 6290, F-35000 Rennes, France

3 – Present address: Laboratory of Biochemistry, Wageningen University & Research, Stippeneng 4, 6708 WE, Wageningen, The Netherlands

#### Supplementary Figures and legends

Fig. S1

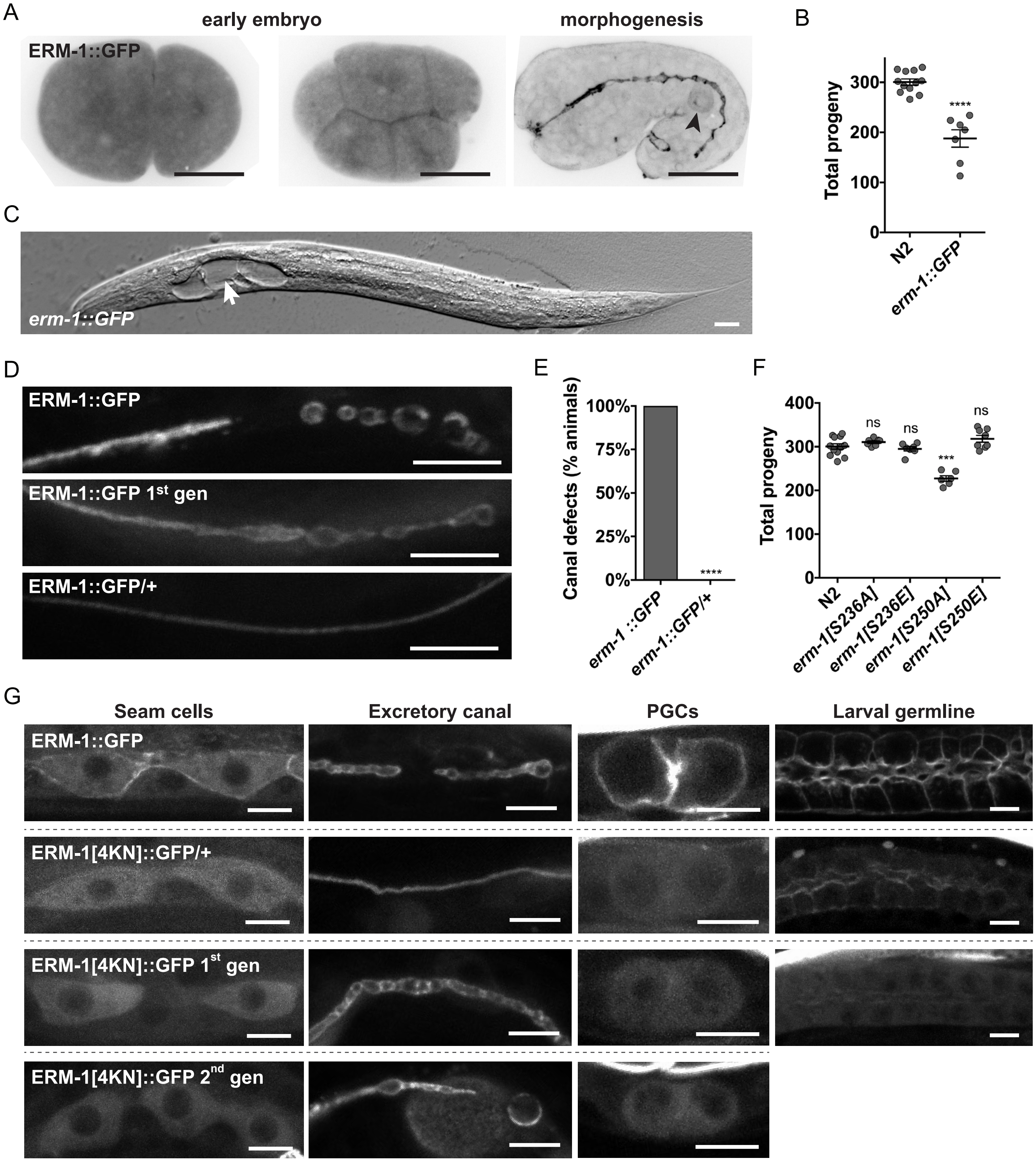
A
early embryo
morphogenesis
ERM-1::GFP
B
Total progeny
300
200
100
0
****
N2
erm-1::GFP
C
erm-1::GFP
D
ERM-1::GFP
ERM-1::GFP 1st gen
ERM-1::GFP/+
E
Canal defects (% animals)
100%
75%
50%
25%
0%
****
erm-1 ::GFP
erm-1::GFP/+
F
Total progeny
400
300
200
100
0
ns
ns
***
ns
N2
erm-1[S236A]
erm-1[S236E]
erm-1[S250A]
erm-1[S250E]
G
Seam cells
Excretory canal
PGCs
Larval germline
ERM-1::GFP
ERM-1[4KN]::GFP/+
ERM-1[4KN]::GFP 1st gen
ERM-1[4KN]::GFP 2nd gen

**Figure S1** - (**A**) Distribution of ERM-1::GFP in different embryonic stages. Arrowhead indicates a germ cell. (**B**) Quantification of total progeny. Each symbol represents the progeny of an individual animal; n≥7. Statistical comparisons are with N2. (**C**) DIC microscopy image of an *erm-1::GFP* L3 larva with severe excretory canal defects. Arrow points to a large cyst close to the excretory canal cell body. (**D**) Excretory canal morphology in ERM-1::GFP animals kept as homozygous for several generations as well as homozygous and heterozygous progeny of ERM-1::GFP heterozygotes for wild type *erm-1*. (**E**) Percentage of animals with defects in excretory canal morphology; n≥40 - Fisher's exact test, comparisons are with ERM-1::GFP. (**F**) Quantification of total progeny. Each symbol represents the progeny of an individual animal; n≥6. Statistical comparisons are with N2. (**G**) Distribution of GFP-tagged wild-type and 4KN ERM-1 variants in different tissues. ERM-1[4KN]::GFP was imaged in 1$^{st}$ and 2$^{nd}$ generation homozygotes and in heterozygous animals. Scale bars: 10 μm; (C) 20 μm; (G) 5 μm.

Fig. S2

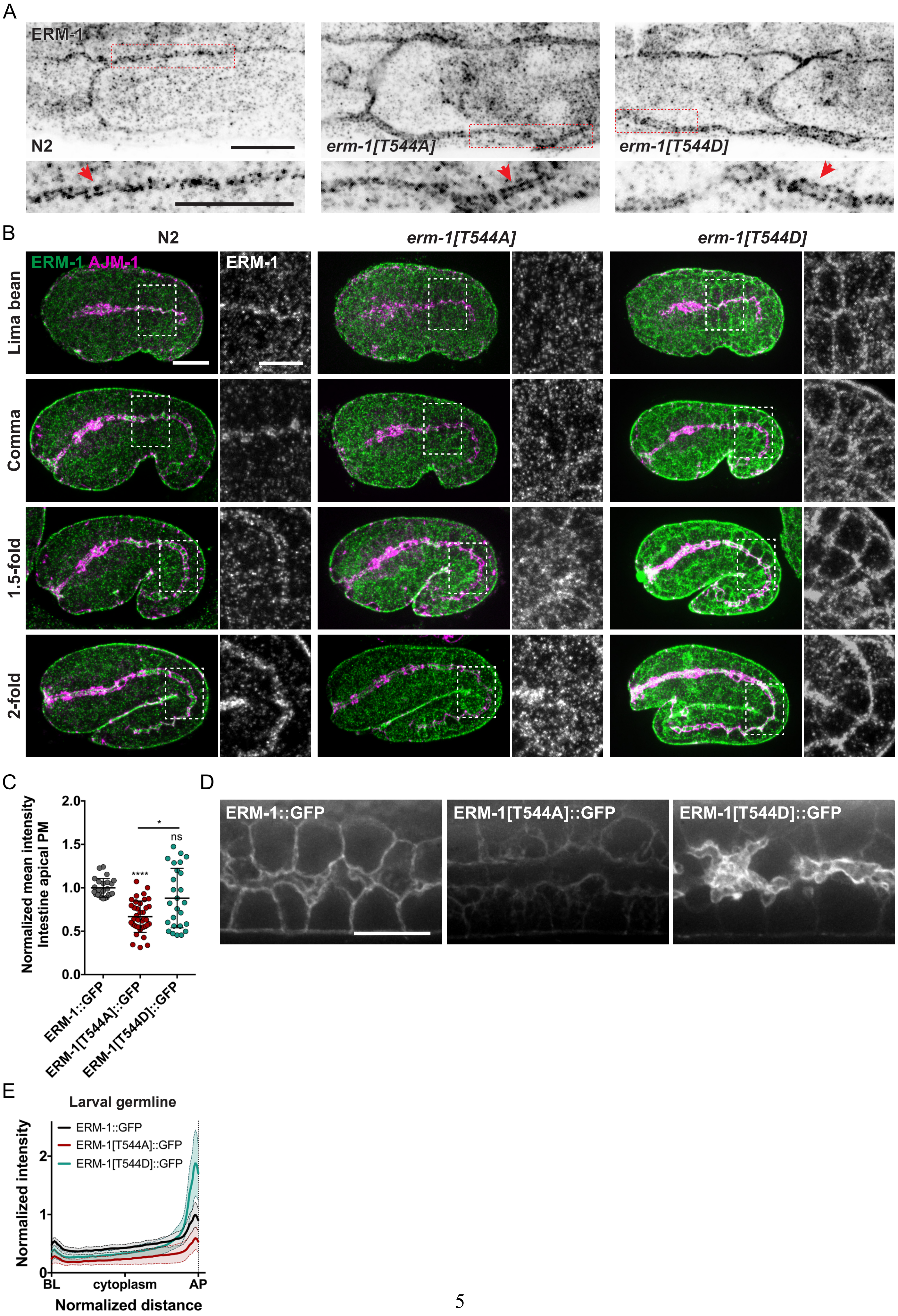
A
ERM-1
N2
erm-1[T544A]
erm-1[T544D]
B
N2
erm-1[T544A]
erm-1[T544D]
ERM-1 AJM-1
ERM-1
Lima bean
Comma
1.5-fold
2-fold
C
Normalized mean intensity
Intestine apical PM
2.0
1.5
1.0
0.5
0.0
****
*
ns
ERM-1::GFP
ERM-1[T544A]::GFP
ERM-1[T544D]::GFP
D
ERM-1::GFP
ERM-1[T544A]::GFP
ERM-1[T544D]::GFP
E
Larval germline
ERM-1::GFP
ERM-1[T544A]::GFP
ERM-1[T544D]::GFP
Normalized intensity
2
1
0
BL
cytoplasm
AP
Normalized distance

**Figure S2 – Distribution of ERM-1 variants in intestine, excretory canal, and germline.** (**A**) Distribution of endogenous ERM-1 in the excretory canal by antibody staining of fixed L2 larvae. Bottom panels are enlarged views of the highlighted regions. Arrows point to the putative apical membrane. (**B**) Distribution of endogenous ERM-1 and AJM-1 by antibody staining in fixed embryos of different stages. Panels to the right are enlarged views of the ERM-1 channel in the highlighted region. (**C**) Mean GFP intensity at the apical membrane in ERM-1::GFP, ERM-1[T544A]::GFP and ERM-1[T544D]::GFP. Each dot represents an individual animal, n≥16. Unless indicated otherwise by a connecting line, statistical comparisons are with ERM-1::GFP. (**D**) Distribution of GFP-tagged ERM-1 variants in the germline of live L4 larvae. Images are maximum intensity projections, and were acquired and displayed with the same settings for comparison. (**E**) Distribution plot of GFP mean fluorescence intensity along the apical-basolateral axis in the germline of L4 larvae. Intensity was normalized to the average peak intensity of wild type animals at the apical membrane, and length was normalized to a fixed distance between the basolateral and apical membranes (see methods; n≥16). Scale bars: 10 μm; enlarged views in (A) 5 μm.

Fig. S3

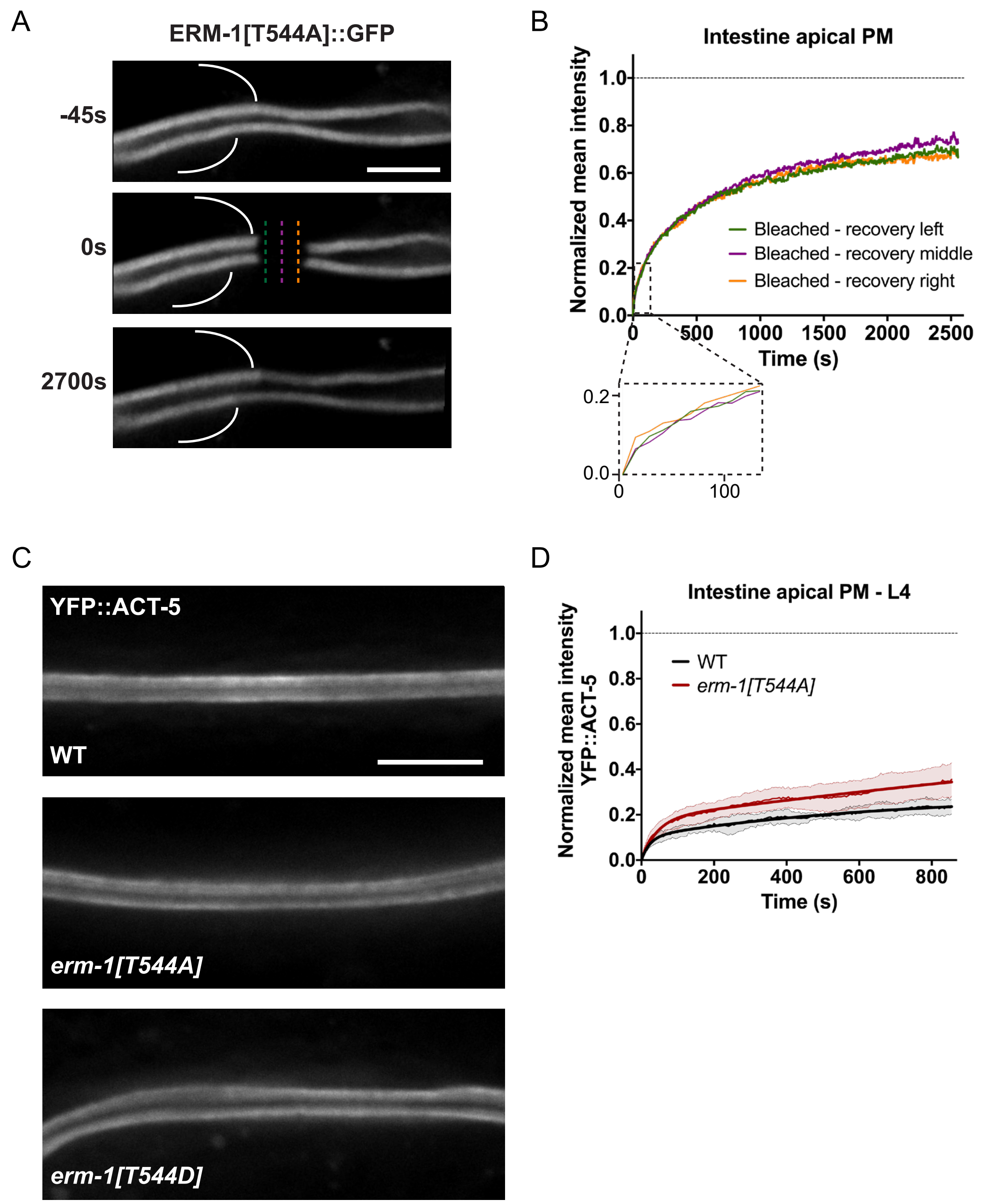
A
ERM-1[T544A]::GFP
-45s
0s
2700s
B
Intestine apical PM
Normalized mean intensity
Bleached - recovery left
Bleached - recovery middle
Bleached - recovery right
Time (s)
C
YFP::ACT-5
WT
erm-1[T544A]
erm-1[T544D]
D
Intestine apical PM - L4
Normalized mean intensity YFP::ACT-5
WT
erm-1[T544A]
Time (s)

**Figure S3** – (**A,B**) Analysis of ERM-1[T544A]::GFP recovery at the apical membrane of L1 larvae intestines. (A) Stills from time-lapse movie. Curved lines indicate lateral membranes separating neighbor and bleached cells. (B) Variation in normalized mean intensity over time in non-bleached areas from neighbor and bleached cells, and in three different positions along the bleached region. Measurements were done in the positions indicated by the color-coded bars in (A). All intensity profiles were corrected for acquisition photobleaching using a separate non-bleached region in neighbor cells. Dashed box below shows a detail of fluorescence recovery within the bleached region in the first 50s. (**C**) Lumen morphology and YFP::ACT-5 distribution in the intestine of L4 larvae. (**D**) FRAP curves of apical YFP::ACT-5 in the intestine of L4 larvae n≥8. Scale bars: 10 μm.

### Supplementary tables

**Supplementary table 1** - Two-component association curve fit analysis of FRAP data (GraphPad Prism 7)

| Tissue | ERM-1 variant | Plateau | Fast half-time (s) | Slow half-time (s) | Goodness of fit R2 |
|---|---|---|---|---|---|
| Intestine AP | ERM-1::GFP | 19% | 8,3 | 1276,0 | 0,991 |
| | ERM-1[T544A]::GFP | 80% | 61,1 | 822,6 | 0,997 |
| | ERM-1[T544D]::GFP | 79% | 17,3 | 986,1 | 0,999 |
| Germline BL | ERM-1::GFP | 73% | 8,4 | 117,2 | 0,992 |
| | ERM-1[T544A]::GFP | 60% | 5,2 | 143,9 | 0,995 |
| | ERM-1[T544D]::GFP | 44% | 12,1 | 260,0 | 0,994 |

**Supplementary table 2** - List of strains used

| Strain | Genotype |
|---|---|
| N2 | wild type |
| GE24 | pha-1(e2123ts) III |
| BOX166 | erm-1(mib11[erm-1[4KN]]) I / dpy-5(e61); unc-29(e403) I |
| BOX213 | erm-1(mib15[erm-1::eGFP]) I |
| BOX233 | erm-1(mib21[erm-1[4KN]::GFP]) I / dpy-5(e61); unc-29(e403) I |
| BOX207 | erm-1(mib14[erm-1[S236A]]) I |
| BOX206 | erm-1(mib13[erm-1[S236E]]) I |
| BOX225 | erm-1(mib20[erm-1[S250A]]) I |
| BOX216 | erm-1(mib17[erm-1[S250E]]) I |
| BOX165 | erm-1(mib10[erm-1[T544A]]) I |
| BOX163 | erm-1(mib9[erm-1[T544D]]) I |
| ML846 | vha-5(mc38) IV; mcEx337[vha-5(+)::GFP; rol-6(su1006)] |
| BOX167 | erm-1(mib10[erm-1[T544A]]) I; vha-5(mc38) IV; mcEx337[vha-5(+)::GFP; rol-6(su1006)] |
| BOX168 | erm-1(mib9[erm-1[T544D]]) I; vha-5(mc38) IV; mcEx337[vha-5(+)::GFP; rol-6(su1006)] |
| N/A | hmr-1(he298[hmr-1::eGFP]) I; dlg-1(mib23[dlg-1::mCherry]) X |
| BOX297 | erm-1(mib9[erm-1[T544D]]) I; hmr-1(he298[hmr-1::eGFP]) I; dlg-1(mib23[dlg-1::mCherry]) X |
| BOX324 | erm-1(mib10[erm-1[T544A]]) I; hmr-1(he298[hmr-1::eGFP]) I; dlg-1(mib23[dlg-1::mCherry]) X |
| MZE1 | unc-119(ed3) III; cbgIs91[pPept-1:PEPT-1::DsRed;unc-119(+)]; cbgIs98[pPept-1:GFP::RAB-11.1;unc-119(+)] |
| BOX183 | erm-1(mib10) I; unc-119(ed3) III; cbgIs91[pPept-1:PEPT-1::DsRed;unc-119(+)]; cbgIs98[pPept-1:GFP::RAB-11.1;unc-119(+)] |
| BOX179 | erm-1(mib9) I; unc-119(ed3) III; cbgIs91[Ppept-1::PEPT-1::dsRED, unc-119(+)]; cbgIs98[Ppept-1::GFP::RAB-11.1, unc-119(+)] |
| BOX218 | erm-1(mib19[erm-1[T544A]::GFP]) I |
| BOX215 | erm-1(mib16[erm-1[T544D]::GFP]) I |
| BOX369 | erm-1(mib19[erm-1[T544A]::GFP]) I; dlg-1(mib23[dlg-1::mCherry]) X |
| BOX261 | mibEx55[Psulp-4::mCherry::ACT-5; Plin-48::GFP] |
| BOX265 | erm-1(mib10[erm-1[T544A]]) I; mibEx55[Psulp-4::mCherry::ACT-5; Plin-48::GFP] |
| BOX266 | erm-1(mib9[erm-1[T544D]]) I; mibEx55[Psulp-4::mCherry::ACT-5; Plin-48::GFP] |
| JM125 | Is[Pges-1::YFP::ACT-5] |
| BOX196 | erm-1(mib10[erm-1[T544A]]) I; Is[Pges-1::YFP::ACT-5] |
| BOX197 | erm-1(mib9[erm-1[T544D]]) I; Is[Pges-1::YFP::ACT-5] |
| BJ49 | kcIs6[Pifb-2::IFB-2::CFP] |
| BOX256 | erm-1(mib9[erm-1[T544D]]) I; kcIs6[Pifb-2::IFB-2::CFP] |

**Supplementary table 3** - List of oligonucleotides used (5' - 3')

| | |
|---|---|
| **Reagents to generate *erm-1[4KN]*** | |
| sgRNA 1 left primer | aattACATGAGCCTTCTTATCAAT |
| sgRNA 1 right primer | aaacATTGATAAGAAGGCTCATgt |
| sgRNA 2 left primer | tcttGATCAAACCAATTGATAAGA |
| sgRNA 2 right primer | aaacTCTTATCAATTGGTTTGATC |
| ssODN repair template | GGATTCCCATGGTCGGAGATTCGTAATATATCATTCAACGACAAtAAtTTTGTCATCAAgCCgATcGAcAAtAAtGCTCATgtaagtgattactgtcacacacagtttgtcgtggcatca |
| integration check left primer | CATCAAgCCgATcGAcAAtAAt |
| integration check right primer | gtagttaccgttgagctgatttg |
| **Reagents to generate erm-1[S236A], erm-1[S236E], erm-1[S250A], and erm-1[S250E]** | |
| sgRNA 1 left primer | tcttGAAAGTCGGATTCCCATGGT |
| sgRNA 1 right primer | aaacACCATGGGAATCCGACTTTC |
| sgRNA 2 left primer | tcttCGCCGAAAGTCGGATTCCCA |
| sgRNA 2 right primer | aaacTGGGAATCCGACTTTCGGCG |
| ssODN repair template S236A | CTTGGATTGAATATTTACGATAAAGCTGATCGTCTTgCGCCGAAgGTtGGtTTtCCgTGGagcGAGATTCGTAATATATCATTCAACGACAAGAAATTTG |
| ssODN repair template S236E | CTTGGATTGAATATTTACGATAAAGCTGATCGTCTTgaGCCGAAgGTtGGtTTtCCgTGGagcGAGATTCGTAATATATCATTCAACGACAAGAAATTTG |
| ssODN repair template S250A | GATTGAATATTTACGATAAAGCTGATCGTCTTTCGCCGAAgGTtGGtTTtCCgTGGagcGAaATcCGaAAcATcgCATTCAACGACAAGAAATTTGTCATCAAACCAATTGATAA |
| ssODN repair template S250E | GATTGAATATTTACGATAAAGCTGATCGTCTTTCGCCGAAgGTtGGtTTtCCgTGGagcGAaATcCGaAAcATcgaATTCAACGACAAGAAATTTGTCATCAAACCAATTGATAA |
| integration check left primer | GAAgGTtGGtTTtCCgTGGagc |
| integration check right primer | GCAGACATCTCctgtacgatc |
| **Reagents to generate erm-1[T544A], erm-1[T544D]** | |
| sgRNA 1 left primer | aattAAGACTCTCCGTCAAATCCG |
| sgRNA 1 right primer | aaacCGGATTTGACGGAGAGTCTT |
| sgRNA 2 left primer | tcttACTCTCCGTCAAATCCGTGG |
| sgRNA 2 right primer | aaacCCACGGATTTGACGGAGAGT |
| ssODN repair template T544A | gcccgattccaaatttttaaattttcagAACAAAAAGGCCGGACGgGAtAAaTAtAAagCTCTgCGaCAgATCCGTGGAGGAAACACAAAACGAAGAATCGATCAATACGAAAAT |
| ssODN repair template T544D | gcccgattccaaatttttaaattttcagAACAAAAAGGCCGGACGgGAtAAaTAtAAagaTCTgCGaCAgATCCGTGGAGGAAACACAAAACGAAGAATCGATCAATACGAAAAT |
| integration check left primer A | GCCGGACGgGAtAAaTAtAAag |
| integration check left primer D | GCCGGACGgGAtAAaTAtAAaga |
| integration check right primer | CCCGAGGAGAAGCACACATG |

| Reagents to generate erm-1::GFP, erm-1[4KN]::GFP, erm-1[T544A]::GFP, erm-1[T544D]::GFP | |
|---|---|
| sgRNA 1 left primer | aattAAGACTCTCCGTCAAATCCG |
| sgRNA 1 right primer | aaacCGGATTTGACGGAGAGTCTT |
| sgRNA 2 left primer | tcttACTCTCCGTCAAATCCGTGG |
| sgRNA 2 right primer | aaacCCACGGATTTGACGGAGAGT |
| LH arm left primer | acgttgtaaaacgacggccagtcgccggcaGGAGgttcgtattttaaaaaaactcg |
| LH arm right primer step 1 | TGATCGATTCTTCGTTTTGTGTTTCCTCCcCtaATcTGcCGGAGAGTCTTGTACTTGTCG |
| LH arm right primer step 1 A | TGATCGATTCTTCGTTTTGTGTTTCCTCCcCtaATcTGcCGcAGAGcCTTGTACTTGTCGCGTCCGG |
| LH arm right primer step 1 D | TGATCGATTCTTCGTTTTGTGTTTCCTCCcCtaATcTGcCGcAGAtcCTTGTACTTGTCGCGTCCGG |
| LH arm right primer step 2 | GATGCTCCTGAGGCTCCCGATGCTCCCATATTTTCGTATTGATCGATTCTTCGTTTTGTG |
| RH arm left primer | CGTGATTACAAGGATGACGATGACAAGAGATAAttatttgttctatcgtatttccttt |
| RH arm right primer | ggaaacagctatgaccatgttatcgatttcgctccatcgaaacccttgga |
| integration check left primer | CTGTCACTGACTACGACGTTCTG |
| integration check right primer | CCCGAGGAGAAGCACACATG |

| Reagents to generate the *Psulp-4::mCherry::ACT-5* construct | |
|---|---|
| Psulp-4 left primer | Cgggccccccctcgaggtcgacggtatcgataagcttgattggattccgcaatgctttga |
| Psulp-4 right primer | tttgaatactggaaaaatagttgc |
| mCherry left primer | cttttaaaataattatgcaactatttttccagtattcaaaATGTCCAAGGGAGAGGAGGA |
| mCherry right primer | CATCGATGCTCCTGAGGCTCCCGATGCTCCCTTGTAGAGCTCGTCCATTCC |
| ACT-5 left primer | GGAGCATCGGGAGCCTCAGGAGCATCGATGGAAGAAGAAATCGCCGCC |
| ACT-5 right primer | tagtggatcccccgggctgcaggaattcgataattatattttaaaaaatgaggggaaataatacacaagt |
